# A genome-scale metabolic reconstruction provides insight into the metabolism of the thermophilic bacterium *Rhodothermus marinus*

**DOI:** 10.1101/2021.05.17.444423

**Authors:** Thordis Kristjansdottir, Gudmundur O. Hreggvidsson, Sigmar Karl Stefansson, Elisabet Eik Gudmundsdottir, Snaedis H. Bjornsdottir, Olafur H. Fridjonsson, Eva Nordberg Karlsson, Justine Vanhalst, Birkir Reynisson, Steinn Gudmundsson

## Abstract

The thermophilic bacterium *Rhodothermus marinus* has mainly been studied for its thermostable enzymes. More recently, the potential of using the species as a cell factory and in biorefinery platforms has been explored, due to the elevated growth temperature, native production of compounds such as carotenoids and EPSs, the ability to grow on a wide range of carbon sources including polysaccharides, and available genetic tools. A comprehensive understanding of the metabolism of production organisms is crucial. Here, we report a genome-scale metabolic model of *R. marinus* DSM 4252^T^. Moreover, the genome of the genetically amenable *R. marinus* ISCaR-493 was sequenced and the analysis of the core genome indicated that the model could be used for both strains. Bioreactor growth data was obtained, used for constraining the model and the predicted and experimental growth rates were compared. The model correctly predicted the growth rates of both strains. During the reconstruction process, different aspects of the *R. marinus* metabolism were reviewed and subsequently, both cell densities and carotenoid production were investigated for strain ISCaR-493 under different growth conditions. Additionally, the *dxs* gene, which was not found in the *R. marinus* genomes, from *Thermus thermophilus* was cloned on a shuttle vector into strain ISCaR-493 resulting in a higher yield of carotenoids.

**Importance:** A biorefinery converting biomass into fuels and value-added chemicals is a sustainable alternative to fossil fuel-based chemical synthesis. *Rhodothermus marinus* is a bacterium that is potentially well suited for biorefineries. It possesses various enzymes that degrade biomass, such as macroalgae and parts of plants (e.g. starch and xylan) and grows at high temperatures (55-77°C) which is beneficial in biorefinery processes. In this study, we reviewed the metabolism of *R. marinus* and constructed a metabolic model. Such a model can be used to predict phenotypes, e.g. growth under different environmental and genetic conditions. We focused specifically on metabolic features that are of interest in biotechnology, including carotenoid pigments which are used in many different industries. We described cultivations of *R. marinus* and the resulting carotenoid production in different growth conditions, which aids in understanding how carotenoid yields can be increased in the bacterium.

## 1. Introduction

*Rhodothermus marinus* is an aerobic bacterium that belongs to the phylum *Rhodothermaeota* [1]. It grows optimally at 65°C on various proteinaceous and carbohydrate substrates and was first isolated from a submarine hot spring in Iceland [2]. The *R. marinus* genome encompasses several gene clusters encoding pathways for utilization and cellular import of diverse carbohydrates. Several enzymes from *R. marinus* have been characterized, and many have biotechnological potential, including a number of polysaccharide degrading enzymes, such as cellulase [3], laminarinase [4] and xylanase [5], [6] (see [7] for a review). *R. marinus* grows on a wide range of sugars from second and third generation biomass and can be cultivated to relatively high yields [8]. It has anabolic pathways and precursor pools for production of various biotechnologically interesting primary and secondary compounds, such as polyamines [9], exopolysaccharides [10], carotenoids [11], compatible solutes [12] and lipids [13], [14].

Anaerobic fermentative organisms are generally preferred to produce low-value commodity chemicals, simple organic acids and alcohols that are typically catabolic waste products from incomplete oxidation of substrates. Conversely, heterotrophic aerobes such as *R. marinus*, typically oxidize their carbon substrates completely under optimized conditions, and therefore utilize organic substrates more efficiently for both energy and carbon. Consequently, aerobes can carry a greater metabolic burden and are the preferred organisms for the anabolic production of complex secondary metabolites.

Cultivation at high temperatures (60-70°C) may be beneficial in bioreactors as it reduces the cost of cooling, and higher temperatures protect the cultures from mesophilic spoilage bacteria. High temperature also increases the solubility of polysaccharides and leads to reduced viscosity of the fermentation broth. This may alleviate scale up problems of mixing and aeration to a significant extent and enable greater substrate loadings. Elevated temperatures may also enable cost-effective recovery of volatile products by distillation or gas stripping, reducing product inhibition and prolonging the production phase of the culture [15].

*R. marinus* has the potential to serve as a robust production organism in the emerging biorefinery industry and as a chassis species that can be metabolically engineered for the production of novel chemical compounds of industrial interest. For this purpose, comprehensive knowledge and understanding of its metabolism are needed. A genome-scale metabolic reconstruction describes the metabolic network of a given organism based on both genomic information and available physiological data. A well-curated reconstruction contains a comprehensive overview of the metabolism of the organism in question. A reconstruction can be converted to a computational model, where phenotypic features are predicted under different conditions. The model can for instance be used to predict growth capabilities on different substrates and guide genetic engineering efforts [16]. Reconstructing the network of a poorly studied organism can result in gaps in the model. While this may skew predictions, it can nevertheless be useful for focusing future research efforts. Although *R. marinus* is not as well studied as common model organisms, it has been the subject of several studies, including the development and application of genetic tools [17], particularly for the engineering of the carotenoid pathway [18].

Carotenoids are pigments produced by many plants, fungi, algae and bacteria. Some non-photosynthetic bacteria produce carotenoids to stabilize cell membrane fluidity in response to extreme environments (high/low temperatures, pH, salinity, etc.) and to protect themselves against UV radiation and oxidative stress [19]. Carotenoids are in demand for different applications, such as the food, feed and cosmetic industries. In a previous study we engineered the carotenoid biosynthetic pathway in *R. marinus* to produce the industrially relevant carotenoid lycopene, instead of native γ-carotenoids [18]. In another, sequential batch cultivation resulted in higher carotenoid production than shake flask cultivation [8]. Drawing upon the metabolic reconstruction of the current study, we further investigated the effects of culture conditions on carotenoid production and growth of *R. marinus*.

Here we reconstructed the genome-scale metabolic network of the *R. marinus* type strain DSM 4252^T^, which has been the subject of most of the published studies so far and for which an annotated genome sequence is available [20]. However, the type strain is not amenable to genetic manipulation as it aggregates in liquid cultures and harbors a highly active DNA restriction enzyme with a 4 base recognition site [21]. Therefore, existing genetic tools were developed for another *R. marinus* strain, ISCaR-493 (DSM 16675), which was selected after screening of numerous *R. marinus* strains for a restriction-deficient phenotype [22]. Here, the genome of strain *R. marinus* ISCaR-493 was sequenced and the genomes of the two strains were compared. This analysis was used to find if any model genes from DSM 4252^T^ could not be found in ISCaR-493, and subsequently if the DSM 4252^T^ model could be extrapolated to strain ISCaR-493. Growth curves and uptake and secretion rates of the main metabolites from bioreactor cultivations were obtained for both strains and the data used to validate the model. During the reconstruction process, several interesting features related to carotenoid production were identified and investigated further. This included heterologous expression of a gene from *Thermus thermophilus* encoding the terpenoid biosynthetic enzyme 1-deoxy D-xylulose 5-phosphate (DXP) synthase, which was not identified in the genomes of *R. marinus*.

## 2. Methods

### 2.1 Strains, media and culture conditions

Three *R. marinus* strains were used in this study, DSM 4252^T^, ISCaR-493 (DSM 16675) and the mutant strain TK-4 (ISCaR-493 derivative, Δ*trpB*Δ*purA::trpBdxs*_*T*.*thermophilus*_, section 2.3). All cultivations were at 65°C and liquid cultures were set to shaking at 200 rpm. For each cultivation, *R. marinus* was first streaked on an agar plate containing rich medium, 10% medium 162 [23], with modifications (2 mM MgSO_4_ and 0.2 mM CaCl_2_ in final volume) and addition of 1% NaCl, 0.03% K_2_HPO_4_, 0.1% yeast extract, 0.1% tryptone, 0.1% peptone, 0.05% glucose, 0.05% starch, 0.06% pyruvate and 0.018% Na_2_CO_3_. Utilization of different carbon sources (section 3.1) was examined on defined medium agar plates containing 10% medium 162 [23] with addition of 8 mM phosphate buffer (KH_2_PO_4_, Na_2_HPO_4_, pH=7.2), 0.1% vitamin solution [23] and 0.4% of each carbon source, except the amino acids which were 0.2%. *R. marinus* strain DSM 4252^T^ from a rich medium agar plate (see above) was resuspended in a drop of 0.9% NaCl solution and subsequently streaked on agar plates containing the different carbon sources. The plates were incubated at 65°C and growth was examined after 1, 3, 5 and 7 days.

The cultivations used to validate the model (section 3.3) were carried out in bioreactors (Labfors 5, Infors HT, Bottmingen, Switzerland), where pO_2_ was kept at 40% with stirrer speed at 200-500 rpm and airflow as needed, pH at 7.2 with addition of 16.5% NH_4_OH and the temperature at 65°C. Defined medium, which contained 10% modified medium 162 with the addition of 1% NaCl, 8 mM phosphate buffer (KH_2_PO_4_, Na_2_HPO_4_, pH=7.2), 10 mM NH_4_Cl, 0.02% asparagine, 0.02% glutamine, 0.1% vitamin solution [23] and the carbon sources 1% glucose and 0.09% pyruvate, was used. The cultures were inoculated with 10% of pre-culture and were performed in duplicates. Samples were taken for OD (620 nm) and HPLC measurements every hour, until a stationary phase was reached (18 – 32h).

The cultivations for examining cell density and carotenoid production under different conditions (section 3.4) were performed in shake flasks under light and dark conditions. All cultures were exposed to day light and additional light from a halogen lamp, except when grown under dark conditions where the flasks were covered from the light. A defined medium (same as in section 3.3, except Wolf’
ss vitamin- and trace elements solution from [24] were used), was used for these cultivations, but with different carbon sources: 1% glucose, 1% glucose and 0.09% pyruvate, 1% alginate, 0.5% glucose and 0.25% pyruvate, 0.09% pyruvate, 0.18% pyruvate, and without a carbon source for a negative control. For cultivation of strain TK-4, adenine (0.0025%) was also supplemented. Two pre-cultures were prepared for each liquid culture to be monitored. First, cells were transferred from a fresh rich medium agar plate (see above) to defined liquid medium containing 0.5% glucose and 0.018% pyruvate and grown overnight (16h). This culture was used to inoculate (10%) a fresh defined liquid medium containing 1% glucose and 0.09% pyruvate, which was also grown overnight (16h). All the monitored cultures were inoculated (10%) with the second pre-culture. The cultures were stopped after 24h and cell density was estimated by measuring OD at 620nm in a spectrophotometer (Novaspec III^+^, Biochrom, Harvard Bioscience Inc., Holliston, Massachusetts, US).

All media components were autoclaved, except for the vitamin solution and the trace element solution, which were filter sterilized. The alginate was not sterilized due to browning and degradation when autoclaved. The probability of contamination in the alginate was low, because of the high growth temperature and because alginate is not a trivial carbon source for most bacteria. The alginate cultures were plated to verify that contamination had not occurred during growth. Individual colonies were obtained, which all had the characteristic red color, and were subsequently identified as *R. marinus* using MALDI-TOF (Microflex, Bruker, Billerica, Massachusetts, US). MALDI-TOF was used according to the manufacturer’
ss instructions.

### 2.2 Analytical methods

The DNA content in *R. marinus* cells was estimated from exponential bioreactor cultures using the fluorometric Quant-iT™ PicoGreen™ dsDNA Assay Kit (ThermoFisher, Waltham, Massachusetts, US). A sample of freeze-dried biomass (approx. 10^7^ cells/ml) was dissolved in water and sonicated for 10 sec to lyse the cells. A standard curve was obtained from PicoGreen measurements of λ-dsDNA and used to estimate DNA concentration in *R. marinus* cells.

Glucose, lactate and acetate concentrations in the samples taken during bioreactor cultivations were measured using high-performance liquid chromatography (HPLC). The samples were filtered through 0.2 µm filters (Phenomenex) and metabolites subsequently quantified using the Dionex 2000 HPLC system (Dionex, Idstein, Germany) with a Rezex ROA-Organic Acid H + (8%, Phenomenex, Aschaffenburg, Germany) and a RI-101 detector (Shodex, München, Germany). Chromeleon evaluation software version 6.80 (Dionex, Idstein, Germany) was used. Separation was obtained using 60°C column temperature with 0.2 mM sulfuric acid (Carl Roth, Karlsruhe, Germany) as the eluent at a flow rate of 600 μl/min for 30 min. External standards of HPLC grade (Merck, Darmstadt, Germany, Sigma-Aldrich, St. Louis, USA) were used. The pyruvate concentration was estimated using a pyruvic acid kit (Megazyme, Bray, Ireland). The two amino acids in the defined medium, asparagine and glutamine, were not measured. Their concentration in the medium was low and did therefore not influence the model predictions to a significant extent. OD (620 nm) values from bioreactor cultivations were converted to CDW (g/l) based on data obtained elsewhere [25], [26] (CDW = 0.75 x OD).

For the carotenoid measurements, the cultures were diluted to OD = 1 at 620 nm. The cell pellets from 1 mL of the diluted cultures were mixed well with 1 mL of acetone and incubated in a sonication bath for 20 min. The samples were centrifuged at 16,000 g for 7 min, resulting in a colorless cell pellet and acetone supernatant containing the extracted carotenoids. The *R. marinus* carotenoids in acetone have maximum absorbance at ∼480 nm (data not shown). The OD at 480 nm of the acetone extracts was measured in a 1 cm quarts cuvette in a spectrophotometer (Novaspec III^+^, Biochrom).

### 2.3 Genetic modification of *R. marinus*

The genomic DNA from *T. thermophilus* HB8 was isolated using the MasterPure Complete DNA purification Kit (Lucigen, Middleton, Wisconsin, US) and used as template for the amplification of the *dxs* (1-deoxy D-xylulose 5-phosphate synthase) gene (TTHA0006) (1848bp) by PCR. The amplification was performed using the Q5 High-Fidelity DNA polymerase (New England BioLabs, Ipswich, Massachusetts, US), according to the manufacturer’
ss instructions. The primers were designed to support HiFi DNA assembly (NEBuilder HiFi DNA Assembly Master Mix, New England BioLabs): dxs_thermus_F (5’-AAACCATGGAGGTGCGCGAT<u>ATGATCTTGGACAAGGTGAAC-3’</u>) and dxs_thermus_R (5’-AGCAGTTCGGTCTCGGCGGTCGACAT<u>TCAGGCCCGTTCATGCACCAAG-3’</u>), with the underlined bases matching the *dxs* gene and the rest matching a *Sal*I (New England BioLabs) digested *R. marinus* shuttle vector pRM3000.0. The pRM3000.0 vector is the pRM3000 vector [17] with an ampicillin resistance gene added, for selection in *E. coli*. The pRM3000.0-*dxs* vector was introduced into chemically competent *E. coli* (NEB 5-alpha) and plated on L-medium [27] with 100 µg/mL ampicillin. After incubation overnight at 37°C, positive clones were identified by amplifying the *dxs* gene, using the *Taq* DNA polymerase (New England BioLabs) according to the manufacturer’s instructions. The primers dxs_thermus_F and R were used. The pRM3000.0*-dxs* vector was isolated from positive *E. coli* clones using the Monarch Plasmid Miniprep Kit (New England BioLabs).

The pRM3000.0-*dxs* vector was introduced into *R. marinus* SB-62 (ISCaR-493 derivative, Δ*trpB*Δ*purA*) by transformation [28], using the *trpB* gene for Trp+ selection. Competent *R. marinus* cells were prepared and transformed as described elsewhere [22]. The GenePulser Xcell electroporation system (Bio-Rad) was used for the transformation, with pulses delivered at 20 kV/cm. For each transformation, 40 µl of washed cells and 1 µg of DNA in ≤ 5 µL volume were used. The pRM3000.0 vector was used as a positive control and sterile MilliQ water as a negative control. Transformed cells were spread on agar plates containing medium 162 [23], with modifications (2 mM MgSO_4_ and 0.2 mM CaCl_2_ in final volume) and additions of 1% NaCl, 0.053% NH_4_Cl, 0.2% soluble starch, 0.2% casamino acids, vitamin solution [23], 0.0025% adenine and 2.5% agar. Plates were incubated at 65°C for 5 days. Positive *R. marinus* clones were verified using the same PCR as positive *E. coli* clones.

### 2.4 Genome sequencing and analysis

*R. marinus* strain ISCaR-493 was cultured in rich liquid medium (see section 2.1). For sequencing by short-read technology, genomic DNA was extracted using the MasterPure Complete DNA purification Kit (Lucigen) and sequencing libraries made by both the Nextera XT (FC-131-1024) and Nextera Mate Pair (FC-132-1001) methods. The two resulting libraries were sequenced on the MiSeq sequencing platform using V3 2x 300bp and V2 2x 250bp chemistry, respectively. The sequence reads were quality assessed using FastQC (v0.11.7) [29] and trimmed using Trimmomatic (v0.39) [30]. A genome was assembled using SPAdes (v3.14.0) [31] with flag for isolates. Genome polishing was done by mapping all shotgun short-reads (Nextera XT) to the largest contig from the *de novo* assembly and generating a consensus sequence using the highest quality bases, using Geneious (v9.1.4). The genome was annotated using the PGAP pipeline from NCBI [32].

### 2.5 Metabolic reconstruction and simulations

A protocol for reconstructing genome-scale metabolic models [33] was used to guide the reconstruction process. A draft reconstruction, based solely on the annotated genome sequence, was obtained using the Model SEED [34]. Since no well-curated model of a closely related bacterium was available at the time of the reconstruction, the obtained draft was neither comprehensive nor functional. It was therefore used as a reference model, along with the well-curated model of *E. coli* (iJO1366) [35]. The *R. marinus* model was manually reconstructed based on these two reference models and *R. marinus* experimental data, obtained here and from the literature. The BiGG [36], BRENDA [37], KEGG [38] and MetaCyc [39] databases and the BLAST tool [40] were extensively used.

Cobrapy [41] was used in all model simulations, along with the GLPK solver. The corresponding code can be found as a Jupyter notebook on Github (https://github.com/steinng/rmarinus). For all simulations, flux balance analysis (FBA) was used [42], [43]. Exchange reactions corresponding to metabolites taken up from the media (glucose and pyruvate) and secreted (lactate and acetate) during growth, were constrained with experimentally obtained rates (Supplementary file 1). FBA was subsequently used to optimize for growth by maximizing flux through the biomass reaction.

For accurate growth rate predictions, the biomass reaction should ideally be based on data obtained for the target organism. Here, the biomass reaction was formulated based mostly on available data on *R. marinus* (Supplementary file 2). Separate biosynthetic reactions for each group of macromolecules (protein, lipid, DNA, etc.) were formulated, describing the ratio of the building blocks (amino acids, fatty acids, nucleotides, etc.) and the energy required. Sensitivity analysis, which shows how much variation in each macromolecule affects the predicted growth rate, was performed. This analysis helps to identify which biomass components most urgently need to be accurately measured.

## 3. Results and discussion

### 3.1 Reconstruction of a genome-scale metabolic model of *R. marinus* DSM 4252^T^

A genome-scale metabolic model of *R. marinus* DSM 4252^T^ was reconstructed, named Rmarinus_578 (https://github.com/steinng/rmarinus). The main features of the reconstruction are listed in Table 1. The Memote tool [44] was used to help guide the reconstruction process, by verifying stoichiometric consistency, mass and charge balance and annotation quality (Supplementary file 3). Reactions and metabolites were usually abbreviated in accordance with the BiGG database and annotations with links to external databases are included. The genome sequence for strain DSM 4252^T^ was obtained from GenBank (accession nr: NC_013501). The genes in the reconstruction were identified with the locus tags from the GenBank file. They were annotated with the old gene locus tag from the GenBank file, the protein ID, protein annotation and protein sequence. Experimental data on *R. marinus* obtained in this study and the available literature was used to curate reactions, genes and gene-protein-reaction (GPR) rules. Several metabolic features were reviewed during the reconstruction process. In the following, we highlight a few, which are of interest for biotechnological application of *R. marinus*.

**Table 1.** Main features of the genome-scale metabolic model of *R. marinus* DSM 4252^T^, Rmarinus_578.

| Genome features |  |
| --- | --- |
| Genome size | 3.26 Mb |
| Protein coding sequences | 2889 |
| Model features |  |
| Genes | 578 |
| Metabolites (unique) | 871 (784) |
| Reactions (with GPR) | 929 (771) |
| Memote total score | 55% |

#### Sugar metabolism

*R. marinus* produces pyruvate from glucose through the Embden-Meyerhof-Parnas (EMP) pathway. A 13C metabolic flux study of the central metabolism in *R. marinus* [45] showed that the EMP pathway and the TCA cycle are both highly active while metabolizing glucose. The oxidative pentose phosphate pathway and the glyoxylate shunt had very low activity and the Entner-Doudoroff (ED) pathway, malic enzyme and phosphoenolpyruvate carboxykinase were inactive.

Growth of *R. marinus* strain DSM 4252^T^ was tested on many different carbon sources, both *in vivo* and *in silico* (Table 2). Growth has been shown on several mono-, di- and polysaccharides, which was also observed *in silico*. However, growth on cellulose was predicted *in silico* while not observed *in vivo. R. marinus* does contain genes encoding a endocellulase (EC 3.2.1.4) [3], [46], which can degrade cellulose into differently sized cellooligosaccharides and cellobiose. However, low specific activities were reported compared to benchmark enzymes. The strain does not contain genes encoding specific exocellulases (EC 3.2.1.91), which degrade glucans into β-cellobioses. However, it does contain several genes encoding glycoside hydrolases, that can degrade different oligosaccharides, including cellobiose and cellooligosaccharides, to monosaccharides [47]. Although *R. marinus* might be able to use some of the cellulose partially degraded by the cellulase and glycoside hydrolases, as the model predicts, the cellulose-specific activity of the enzymes is probably not high enough for it to grow solely on cellulose. *R. marinus* was unable to grow *in vivo* on several of the tested carbon sources, especially organic- and amino acids (Table 2). The model predictions usually showed the same results. However, when transport reactions were added to the model, growth was often observed *in silico* for carbon sources that did not result in growth *in vivo*. These carbon sources might be good targets for adaptive evolution experiments. Promiscuous enzymes might adapt to transport these compounds into the cell, where they can presumably be used for growth [48].

**Table 2:**
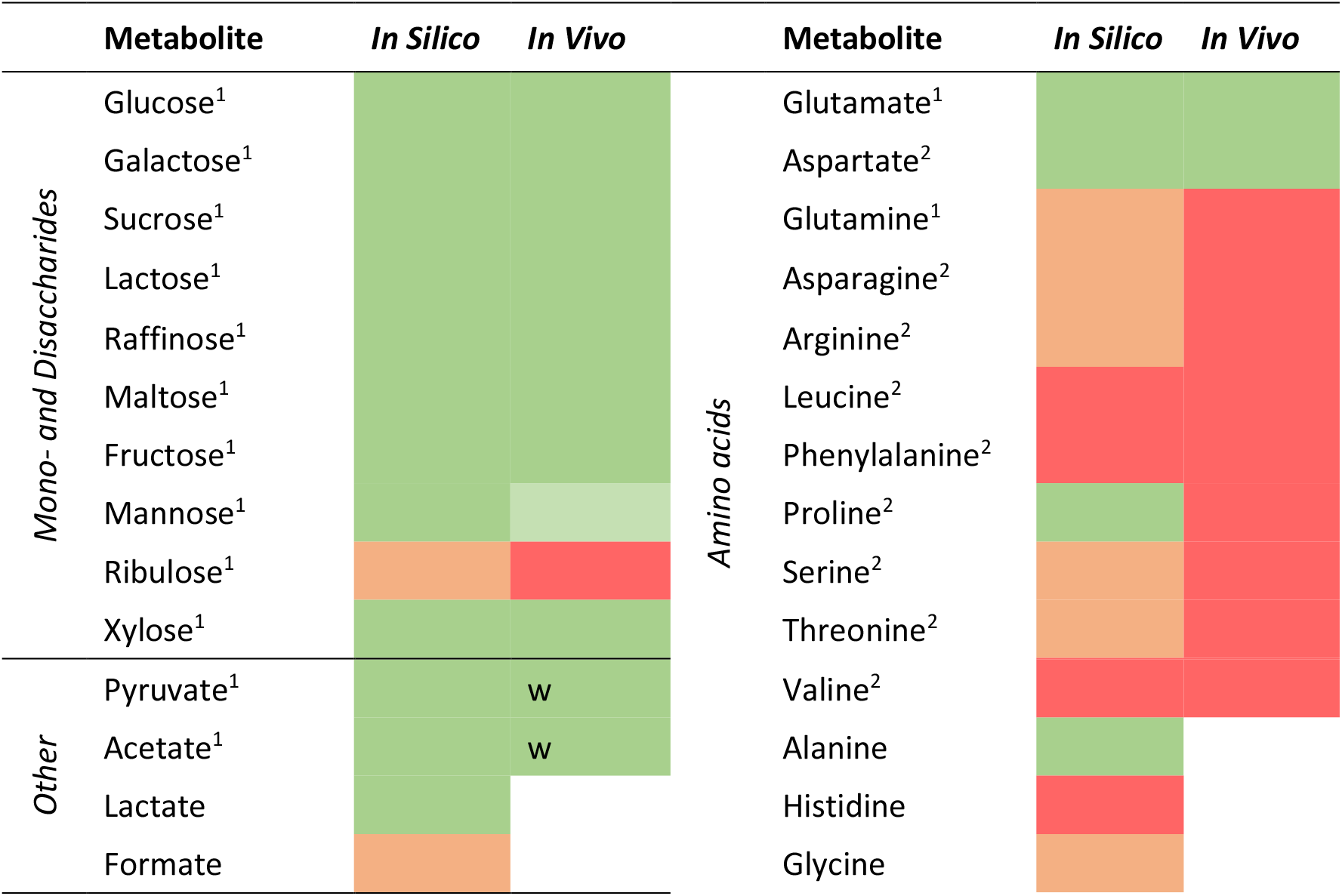
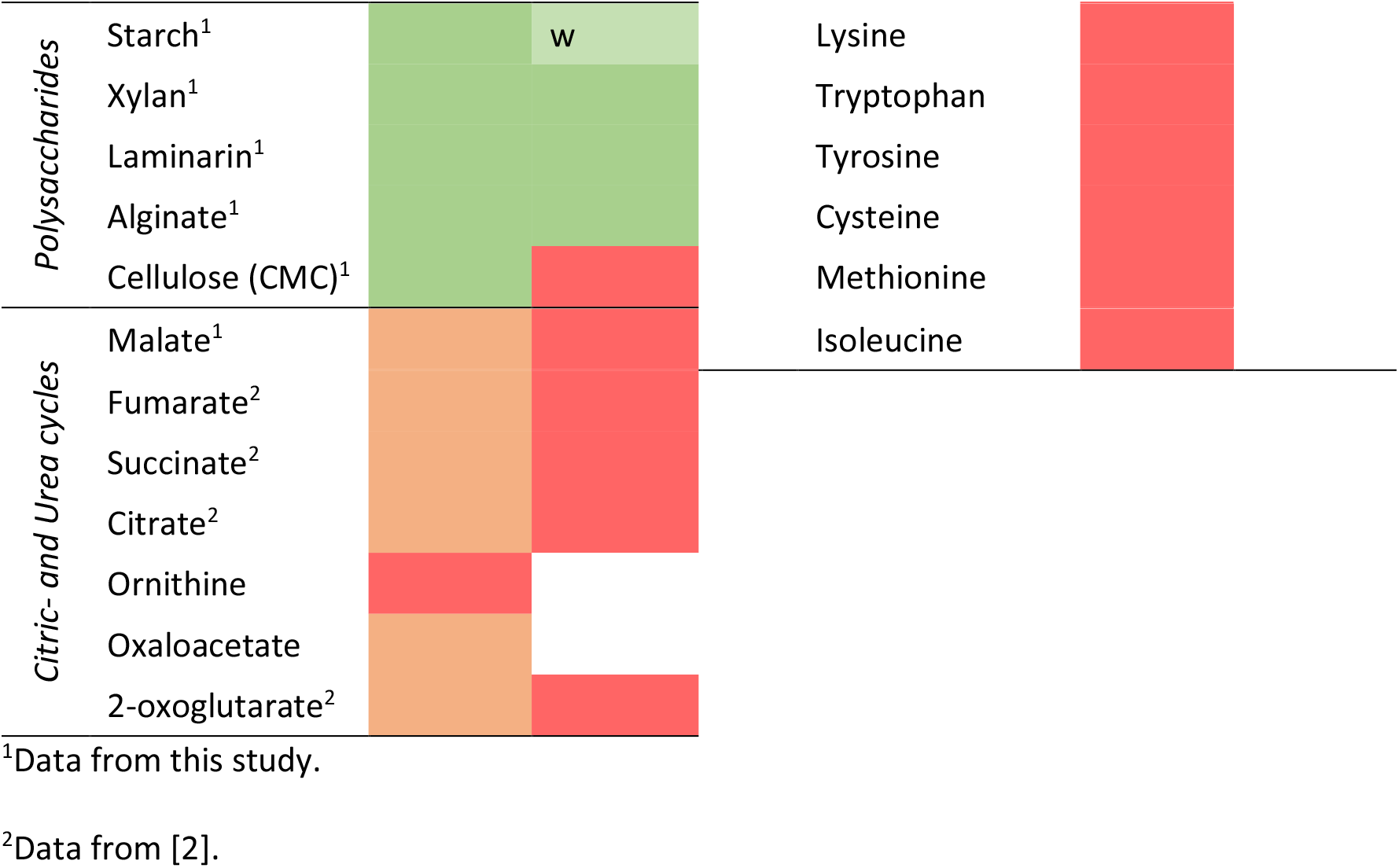
Growth of *R. marinus* strain DSM 4252^T^ on different carbon sources investigated *in silico* and *in vivo*. Green indicates growth, light green indicates weak growth, ‘w’ indicates white colonies (instead of the characteristic red), orange indicates no growth *in silico* except if a transport reaction was added to the model, red indicates no growth and no color indicates no data available.

Lactose and galactose are examples of the di- and monosaccharides that *R. marinus* can use for growth (Table 2). The gene encoding a β-galactosidase (EC 3.2.1.23), which hydrolyzes lactose into glucose and galactose, was found in the genome. Three steps are needed to convert galactose to glucose-6-phosphate, which then enters the glycolysis EMP pathway. The genes encoding galactokinase and galactose-1-phosphate uridylyltransferase, catalyzing the first two steps (galactose -> galactose-1-phosphate -> glucose-1-phosphate), were found in the genome. The third step, where glucose-1-phosphate is turned into glucose-6-phosphate, is usually performed by the enzyme phosphoglucomutase (EC 5.4.2.2). The gene for this enzyme was not found in the genome. However, a homology search showed similarity between known phosphoglucomutase genes from other bacteria and genes RMAR_RS01880 (E value 2e-37) and RMAR_RS08875 (E value 1e-25) which are annotated as phosphomannomutase (EC 5.4.2.8) and phosphoglucosamine mutase (EC 5.4.2.10), respectively. The enzyme phosphoglucosamine mutase in *E. coli*, which usually catalyzes the interconversion of glucosamine-6-phosphate and glucosamine-1-phosphate, was also shown to be able to catalyze the interconversion of glucose-6-phosphate and glucose-1-phosphate, at a lower rate [49]. The phosphorylation site of this enzyme in *E. coli* is Ser102 and a mutational change of Ser100 to a threonine residue increased the phosphoglucomutase activity significantly. Gene RMAR_RS08875 from *R. marinus* was investigated and the serine residue responsible for the phosphorylation was found to be residue number 103. The corresponding residue to Ser100 in the *E. coli* enzyme was found to be a threonine, which indicates that this *R. marinus* enzyme may be responsible for the interconversion of glucose-6-phosphate and glucose-1-phosphate in *R. marinus*.

*R. marinus* possesses several genes encoding polysaccharide degrading enzymes. As a marine bacterium, seaweed is common in its natural environment. Alginate and laminarin are major polysaccharides of brown algae, which *R. marinus* can break down and use as sole carbon sources for growth [50]. Alginate is a structural component of brown algae and can comprise up to 40% of its dry matter [51]. It is a polyuronate that consists of β-D-mannuronate (M) and α-L-guluronate (G) units forming (1 → 4) linked G-, M- and mixed blocks in the polysaccharide chain. The *R. marinus* genome has four genes encoding alginate lyases [52], [53] that, together, depolymerize alginate into the same unsaturated mono-uronate derivative of the M and G units. The *R. marinus* genome also possesses the genes encoding for the remaining enzymes of the alginate catabolic pathway enabling its utilization. The unsaturated monouronate is converted to 4-deoxy l-erythro 5-hexoseulose uronic acid (DEH) y a spontaneous reaction and further catalyzed to 2-keto 3-deoxygluconate (KDG) by an aldose reductase [54]. KDG enters the partial ED pathway in *R. marinus*, where it is catalyzed to 2-keto-3-deoxygluconate 6-phosphate (KDPG) by 2-keto 3-deoxygluconokinase (EC 2.7.1.45) and then by 2-dehydro-3-deoxyphosphogluconate aldolase (EC 4.1.2.14), to pyruvate and glyceraldehyde 3-phosphate which enter central metabolism [54], [55] (Figure 1). *R. marinus* does not possess a key enzyme in the ED pathway, phosphogluconate dehydrogenase (EC 4.2.1.12), which is needed to metabolize glucose. This explains why the ED pathway is not active when *R. marinus* is grown on glucose [45], while the partial pathway is essential for utilization of alginate (Figure 1).

**Figure 1.**
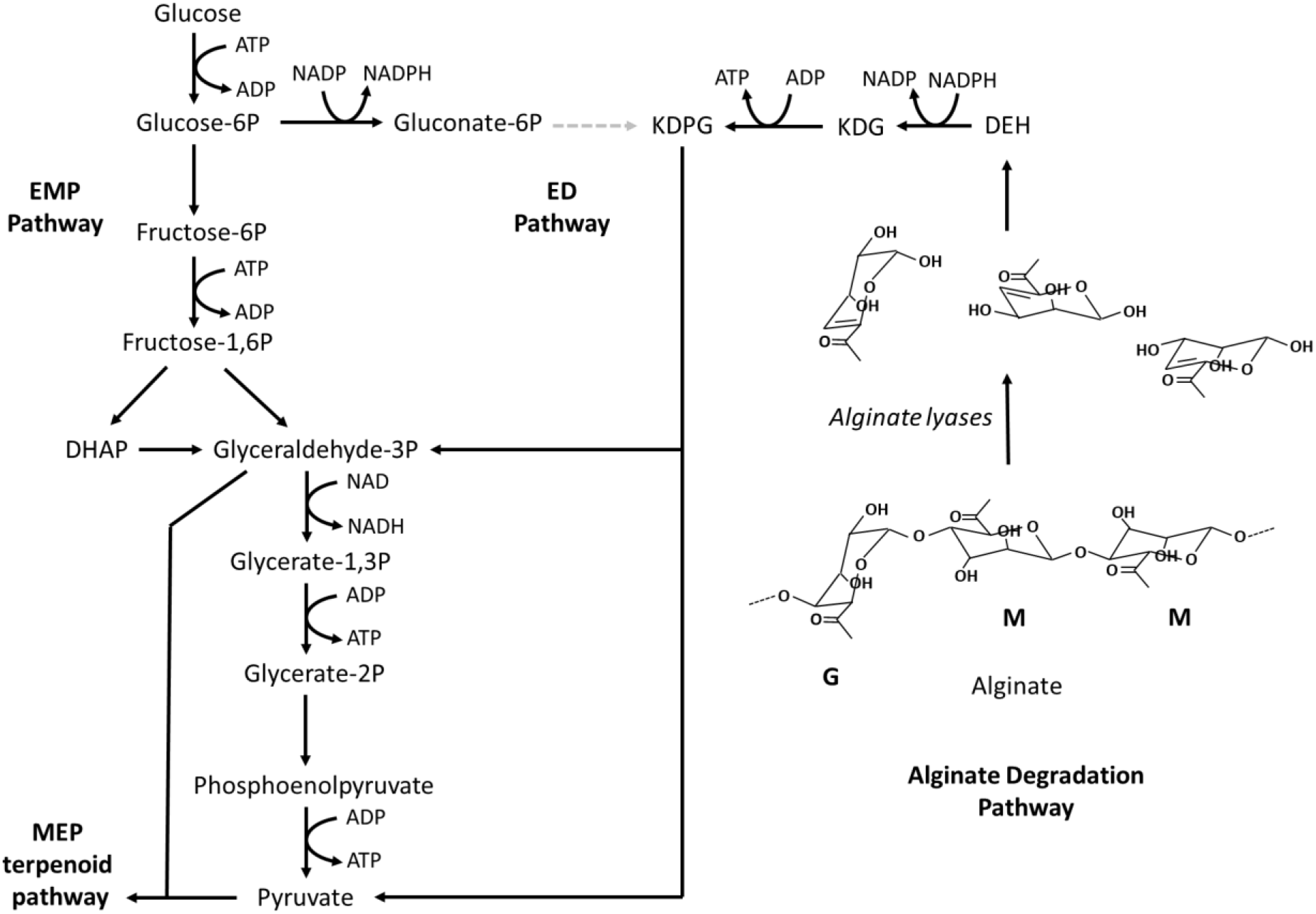
The alginate degradation pathway of *R. marinus* and its connections to the Embden-Meyerhof-Parnas (EMP) pathway, the partial Entner-Doudoroff (ED) pathway and the MEP terpenoid pathway. Names of substrates, products and energy molecules (ATP, NADH, NADPH) are shown, some abbreviated: 4-deoxy-L-erythro 5-hexoseulose uronic acid (DEH), 2-keto 3-deoxygluconate (KDG) and 2-keto 3-deoxygluconate 6-phosphate (KDPG). The molecular structure of a partial alginate molecule is shown: β-D-mannuronate (M) and α-L-guluronate (G). The missing reaction of the ED pathway in *R. marinus* is represented by a grey dotted arrow.

#### Terpenoid and carotenoid metabolism

The carotenoids produced by several *R. marinus* strains have been characterized [11], [56]. A carotenoid pathway was proposed in [11] using structural and bioinformatic data, and later refined [18]. This pathway is included in the reconstruction along with candidate genes. Terpenoids, which serve as precursors to carotenoids, can be produced through two different pathways, the mevalonate [57], [58] and the non-mevalonate (MEP) [59] pathways. In the latter, which is used by most bacteria including *R. marinus*, geranylgeranyl diphosphate (GGDP) is produced from pyruvate and glyceraldehyde 3-phosphate in multiple steps. *R. marinus* possesses genes coding for all the enzymes in the MEP pathway, except for 1-deoxy D-xylulose 5-phosphate (DXP) synthase (DXS, EC 2.2.1.7), which catalyzes the first step in the pathway. Studies of the MEP pathway in other bacteria have shown that DXS is not strictly necessary for the synthesis of DXP. Examples include a mutated form of pyruvate dehydrogenase that is known to rescue *E. coli* cells defective in DXS [60] and a mutated RibB protein and a YajO protein that synthesized DXP from ribulose 5-phosphate, also in *E. coli* [61]. Another possibility to bypass DXS is via the MTA-isoprenoid shunt, as has been shown in *Rhodospirillum rubrum* [62]. Here, the dead-end metabolite of polyamine biosynthesis, 5-methylthioadenosine (MTA) is metabolized by an alternative methionine salvage pathway, which produces DXP as a side-product. Phylogenetic analysis showed that the genes in this pathway are partially present in the *R. marinus* genome. At present, it is not known how DXP is produced in *R. marinus* and without further evidence of alternative pathways, the DXP synthase reaction is present in the reconstruction without any gene candidates assigned. The absence of DXS in *R. marinus* directly suggests heterologous expression of a thermostable DXS as means to increase flux through the terpenoid pathway.

Light-inducible carotenoid production has been observed in many organisms, including non-photosynthetic bacteria, and the regulatory mechanisms have been studied in some of them, including *Myxococcus xanthus* [63], *Thermus thermophilus* [64], *Streptomyces coelicolor* [65] and *Bacillus megaterium* [66]. The MerR family transcriptional regulator, LitR, acts as a repressor in the carotenoid gene cluster. Its activity is dependent on the binding with adenosyl B_12_ and the LitR-AdoB_12_ complex becomes inactivated when illuminated. This causes cell cultures to become colorless under dark conditions while producing carotenoids in light. A homologue for the *T. thermophilus litR* gene was previously identified in the carotenoid gene cluster in *R. marinus* [18], located upstream of the carotenoid gene *crtB* (phytoene synthase). This suggests that light might help increase carotenoid yields in *R. marinus*.

#### Other pathways

The respiratory chain in *R. marinus* has been extensively studied. The first two complexes, NADH dehydrogenase and succinate dehydrogenase, have been characterized [67], [68] and found to be similar to those of other bacteria. The third complex, cytochrome dehydrogenase, is not the typical *bc*_*1*_ but an alternative complex [69]. It has an entirely different structure but carries out the same function, oxidizing reduced menaquinones-7 [70] and reducing High Potential iron-sulfur Protein (HiPIP) and cytochrome *c*. Finally, three different types of the fourth complex have been characterized in *R*. marinus, *cbb3* [71], *caa3* [72] and *ba3* [73]. These reactions and associated genes are included in the reconstruction.

Polyamines are alkaline organic compounds with at least two primary amino groups. They can be found in most forms of life and have diverse functions. Some polyamines are biotechnologically interesting because they can be used to produce plastics and are used as curing agents in polymer applications [74]. In thermophilic bacteria unusual, long and branched polyamines have been observed [75]. They are believed to have protective effects on nucleic acids and proteins under high-temperature conditions. Seven different polyamines have been characterized in *R. marinus* to date, putrescine, spermidine, cadaverine, spermine, thermopentamine, N4-aminopropylspermidine and N4-bis(aminopropyl)spermidine [9], and their biosynthesis is included in the reconstruction. *R. marinus* produces compatible solutes to protect the cell against sudden osmotic changes. They include amino acids, monosaccharides such as trehalose, small peptides and, most abundantly, mannosylglycerate [76]. Mannosylglycerate in *R. marinus* is synthesized via two pathways [77], which have been studied in detail, along with the corresponding enzymes and genes [14], [77], [78]. These pathways are present in the metabolic reconstruction, but as they are produced in response to stress, they are not included in the biomass reaction and the pathways therefore not active during growth simulations where the biomass is maximized.

Genomic information indicates that *R. marinus* can synthesize all the amino acids needed for protein synthesis. This is supported by growth experiments which show that *R. marinus* can grow in defined medium, without any addition of amino acids [79]. Biosynthetic pathways for all the 20 amino acids are included in the reconstruction. The fatty acid and lipid composition of *R. marinus* have been characterized [13], [14]. The dominating fatty acids are iso- and anteiso-C15 and iso- and anteiso-C17, with iso-C16 and iso-C18 as minor components. The major polar lipids are phosphatydilethanolamine, diphosphatydilglycerol and one unidentified lipid and phosphatidylglycerol was identified as a minor lipid [14], [70]. The biosynthesis of the fatty acids and lipids are also included in the reconstruction.

#### The biomass objective function

In genome-scale metabolic models a biomass objective function (BOF) is used to simulate growth. The BOF describes the ratio between the macromolecules (protein, DNA, RNA, lipids, etc.), the composition of each macromolecule (amino acids, nucleotides, fatty acids, etc.) and the energy that the cell requires to grow and maintain itself. We collected data on several studies that describe different components of the biomass in *R. marinus*. Protein, RNA, lipid and glycogen content and amino acid ratio in the biomass were based on [45], carotenoid composition on [56], EPS on [10], lipids on [13], [14] and polyamines on [9]. Quantification of DNA in the biomass was measured in this study (Supplementary file 2). Nucleotide compositions of DNA and RNA were derived from the DSM 4252^T^ genome. The RNA estimate assumes equal transcription of all genes and is therefore expected to be somewhat inaccurate. Growth associated maintenance (GAM) was estimated using experimental data obtained here (supplementary file 1). The remainder of the biomass components was adopted from *E. coli* [80]. A detailed overview of the biomass, sources of information and calculations can be found in supplementary file 2.

Sensitivity analysis was performed to investigate how sensitive the predicted growth rate was to changes in biomass and energy components. The growth rate was predicted while varying each component by 50%, for multiple glucose uptake rates. The components tested were protein, DNA, RNA, lipid, exopolysaccharide, lipopolysaccharide, glycogen, peptidoglycan, GAM and non-growth associated maintenance (NGAM) (Supplementary file 2). The predicted growth rate was most sensitive to changes in the protein component, followed by the lipid component.

### 3.2 Genome comparison of strains DSM 4252^T^ and ISCaR-493

The metabolic model was reconstructed based on genomic information from *R. marinus* DSM 4252^T^. This strain is, however, not amenable to genetic manipulation as it aggregates in liquid cultures and shows high DNA degrading activity due to the presence of a 4 cutter restriction enzyme, *Rma*I [21]. Therefore, genetic tools were developed for a derivative of the strain ISCaR-493 [22]. The 16S rRNA sequence similarity between the two strains is 97.92%.

#### Core genome

The genome of ISCaR-493 was sequenced using Illumina MiSeq and assembled using SPAdes. Subsequently, the 60 resulting contigs were annotated using the NCBI annotation software PGAP. The resulting draft genome was compared to the genome of strain DSM 4252^T^ using the in-house pan-genomic software Genset. The genome of ISCaR-493 had a calculated G+C% of 64.6% compared to 64.3% for strain DSM 4252^T^. The Average Nucleotide Similarity (ANI value) [81] was 95.5%, indicating a relatively high sequence divergence while the genome metrics were very similar for the two strains (Supplementary file 4).

The number of total genes was 2890 and 2937 for ISCaR-493 and DSM 4252^T^, respectively and protein-coding genes were about 98.3% thereof for each strain. 2609 protein-coding genes belonged to the common core having 50% identity across at least 50% of the protein. 74% of the core protein-coding genes could be assigned to COG functional categories [82]. The remaining protein-coding core genes (26%) did not get COG IDs and had unknown (S) or poorly characterized functions (R). 230 protein-coding genes were unique to ISCaR-493 and 275 genes were unique to DSM 4252^T^, a total of 505 genes that comprised the peripheric gene fraction. About 50% of the protein-coding genes in the peripheric fractions of both strains could not be assigned function compared with approximately 26% in the common core (Figure 2).

**Figure 2.**
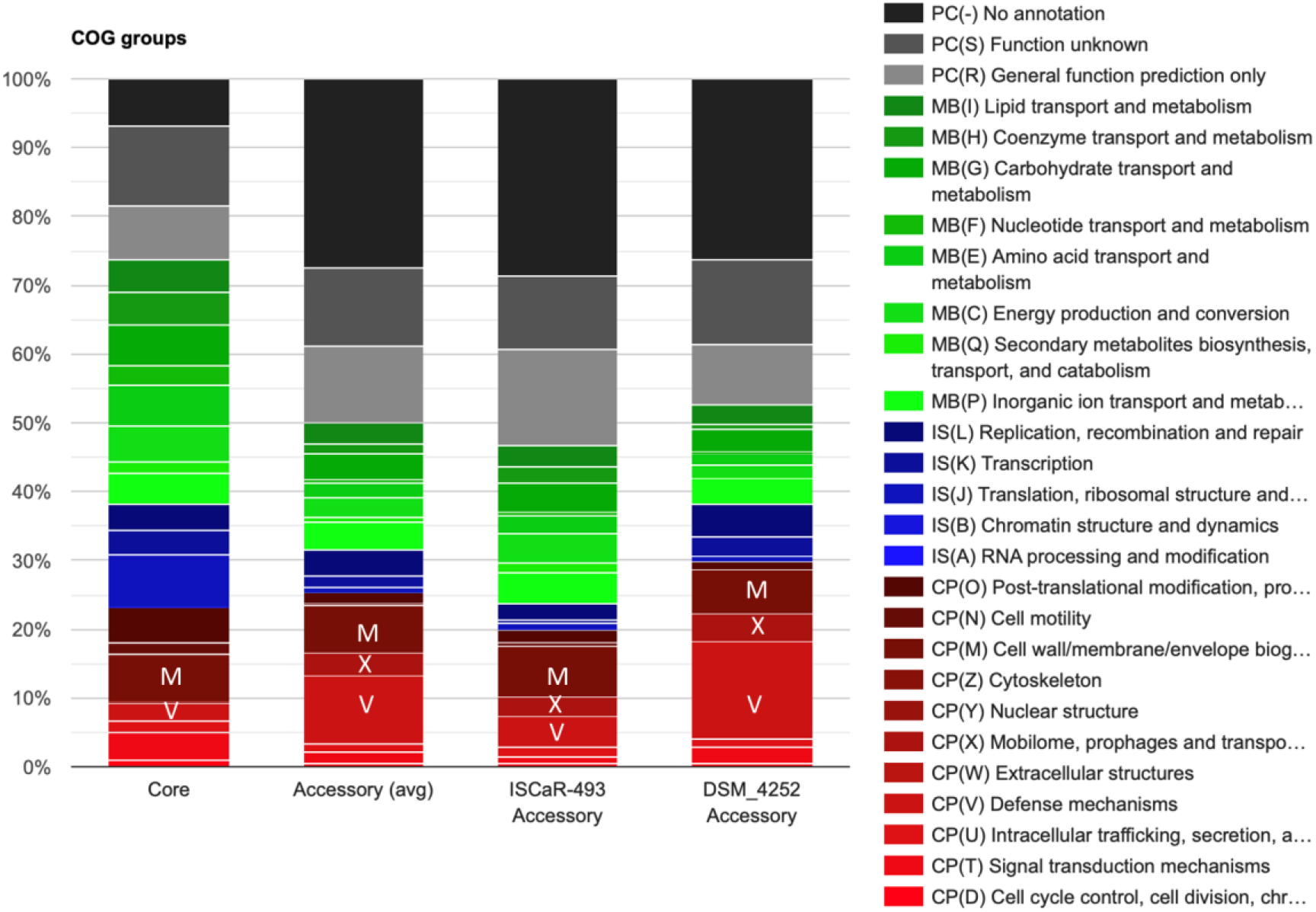
Functional annotations of core and accessory genes in the *R. marinus* strains ISCIaR-493 and DSM 4252^T^. The main functional categories are four: Cellular processes and signalling (CP) red stacks; Information storage and processing (IS) blue stacks; (MB) green stacks; poorly characterized (PC), grey stacks: Single letters designate the COG subcategories. The core genome consists of the predicted genes that are common in both strains. Each category or subcategory is graphed as a percentage of the total number of genes in the core or accessory genomes.

#### Accessory genome

Compared with the core genome the peripheric fractions of both strains were proportionally enriched in the following COG subcategories: V (Defense), M (Cell wall/membrane/envelop biogenesis), and X (Mobilome: Prophages and transposons) (Figure 2). Two restriction enzymes were shown to be only in the peripheric genome of DSM 4252^T^, including the *Rma*l 4 cutter. This reflects the different restriction phenotypes observed [21] and the probably explains the facilitated uptake of foreign DNA in ISCaR-493 compared to DSM 4252^T^.

A total of 578 genes were used in the metabolic network modelling of *R. marinus* 4252^T^ and the great majority of them belonged to the core genome. Only 7 model genes were not found in the genome of ISCaR-493 and only three of them did not have isozymes in the genome that showed high similarity to DSM 4252^T^ genes (Table 3).

**Table 3.** The model genes that are not present in *R. marinus* strain ISCaR-493. The model gene IDs from DSM 4252^T^ are listed, with corresponding annotations and the gene IDs of the isozymes, when applicable.

| DSM 4252<sup>T</sup> gene ID | Annotation | Isozyme present |
| --- | --- | --- |
| RMAR_RS02620 | Dihydroneopterin triphosphate pyrophosphatase | Yes (RMAR_RS06180) |
| RMAR_RS14780 | O-antigen polymerase (wzy) | No |
| RMAR_RS05305 | Xylanase | No |
| RMAR_RS14585 | Alginate lyase | No |
| RMAR_RS11910 | Asparagine synthase (glutamine-hydrolyzing) | Yes (RMAR_RS10310) |
| RMAR_RS12230 | Beta-ketoacyl-ACP synthase | Yes (RMAR_RS14045) |
| RMAR_RS12885 | N-acetylglutamate synthase | Yes (RMAR_RS13420) |

Strain DSM 4252^T^ contains two genes encoding xylanases while strain ISCaR-493 contains only one homologue. Both strains grow on xylan as the sole carbon source (Table 2 for DSM 4252^T^, data not shown for ISCaR-493). Strain DSM 4252^T^ contains four genes encoding alginate lyases and one of them is missing in ISCaR-493. The latter strain grows well in a medium with alginate as the sole carbon source [50], suggesting that the three alginate lyases are sufficient to degrade alginate for utilization.

Many enzymes take part in EPS (the envelope polysaccharides) biosynthesis and assembly, and their corresponding genes were all found in the core genome. A putative o-antigen polymerase (wzy) found in the accessory genome of strain ISCaR-493 showed a low similarity to a functionally corresponding protein in DSM 4252^T^ (E-value 0.004). However, the encoded gene showed high similarity to genes in more distantly related bacteria annotated as o-antigen ligase and o-antigen polymerase. This enzyme activity is essential for EPS synthesis and must be present in ISCaR-493 as EPS is produced [10]. The relatively high content of genes in the accessory genomes belonging to subcategory M (Cell wall/membrane/envelop biogenesis), including glycosyl transferases and sulfotransferases, indicates that the EPS of the two strains may be different in sugar composition and sulfatation patterns and this may explain the differently aggregating phenotypes of the strains.

The above analysis suggests that the Rmarinus_578 model can be used for both strains DSM 4252^T^ and ISCaR-493.

### 3.3 Model validation for strains DSM 4252^T^ and ISCaR-493

Experimental growth data was obtained for strains DSM 4252^T^ and ISCaR-493 in bioreactors, along with measured uptake of glucose and pyruvate and secretion of acetate and lactate (Supplementary file 1). The average rates of two replicates (Supplementary file 1) was used to constrain the model to validate the accuracy of growth predictions (Figure 3a). The experimental growth data showed that the growth of *R. marinus* DSM 4252^T^ did not follow the typical batch growth curve of bacteria. A true exponential phase was not observed throughout the growth phase. Instead, the apparent specific growth rate decreased over time until stationary phase was reached (Figure 3b). The specific growth rate in strain ISCaR-493 was closer to being constant, with exponential growth during a longer period (Figure 3b). Data from time points 3-6 for strain DSM 4252^T^ and 1-5 for strain ISCaR-493 was used here. This analysis showed that the model accurately predicts growth for both strains (Figure 3a).

**Figure 3.**
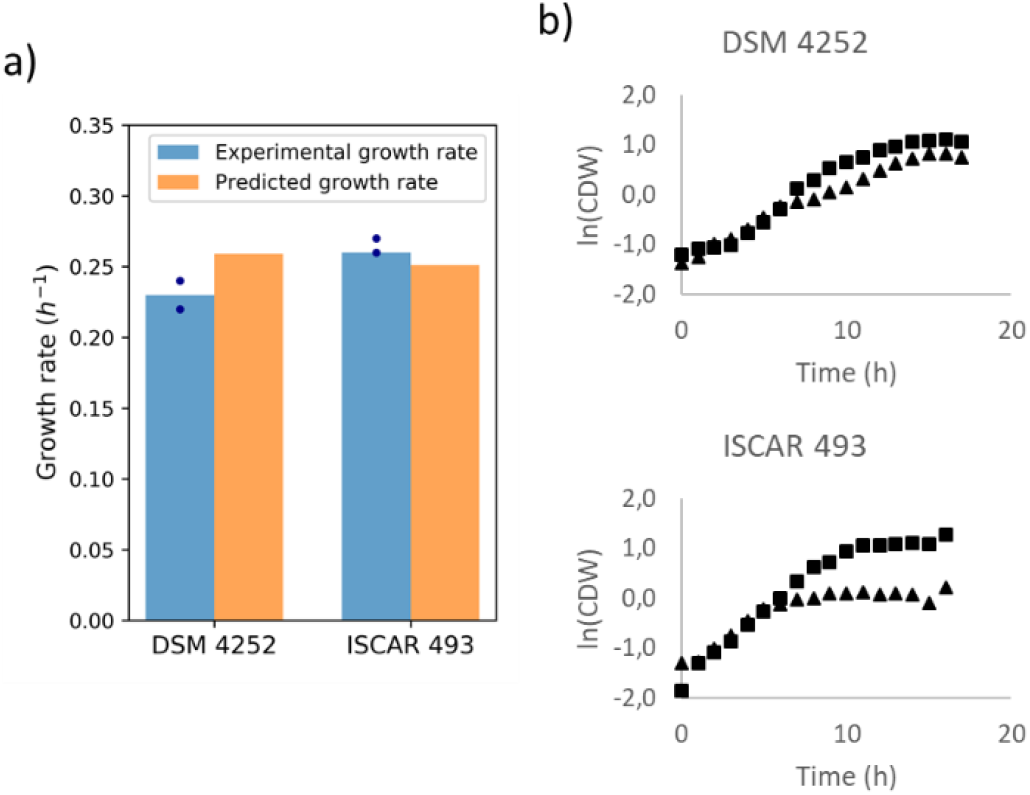
Experimental growth data for strains DSM 4252^T^ and ISCaR-493 was used to validate model growth predictions. The model was constrained with uptake (glucose and pyruvate) and secretion (lactate and acetate) rates observed *in vivo* (Supplementary file 1) and optimized for growth. Predicted growth rates were compared to growth rates observed *in vivo* (**a**). Experimental data from the exponential growth phase of two replicates (▲ and ■), obtained early in the growth phase (time points 3-6 for strain DSM 4252 and 1-5 for strain ISCaR-493), was used (**b**).

For both strains, but more so for DSM 4252^T^, secretion rates of lactate and acetate increased during the growth phase (Supplementary file 1). A decrease in growth rate during batch cultivations has been observed in other bacteria, such as *E. coli* [83] where the main reason was oxygen limitation that could also lead to an increase in organic acid secretion. The cultivations here were carried out with high aeration as oxygen levels were kept fixed at 40% pO_2_. A plausible explanation for why the cells would experience oxygen limitation in a medium with excess oxygen levels is local limitation due to cell aggregation [84]. Aggregation of several *R. marinus* strains has been reported previously [22], especially in DSM 4252^T^ and *R. marinus* is also shown to produce exopolysaccharides [10], which can cause cells to aggregate [85].

When the model was optimized for growth, without oxygen limitation and free secretion of acids, it did not predict any acid production and the predicted growth rate was slightly higher than observed *in vivo*. When oxygen was limited in the model, the predicted growth rate decreased, and the model predicted lactate secretion (data not shown). Experimental data showed that lactate was first secreted, followed by acetate (Supplementary file 1). The model predicted slightly higher growth rate when lactate was the sole acid produced, opposed to when it was forced to also produce acetate.

### 3.4 Carotenoid production and growth of *R. marinus* ISCaR-493

To better understand the carotenoid production in *R. marinus*, a cultivation experiment comparing different conditions was performed. Besides obtaining high yields of carotenoids per cell, high cell density is important for achieving high yields of carotenoids. Therefore, both extracted carotenoids (from 1 mL of cells diluted to OD620 nm = 1) and cell densities were measured from cultivations after 24 hours (Figure 4). The ISCaR-493 strain was used in this experiment, as it can be genetically modified and thus likely to be used for future cell factory designs

**Figure 4.**
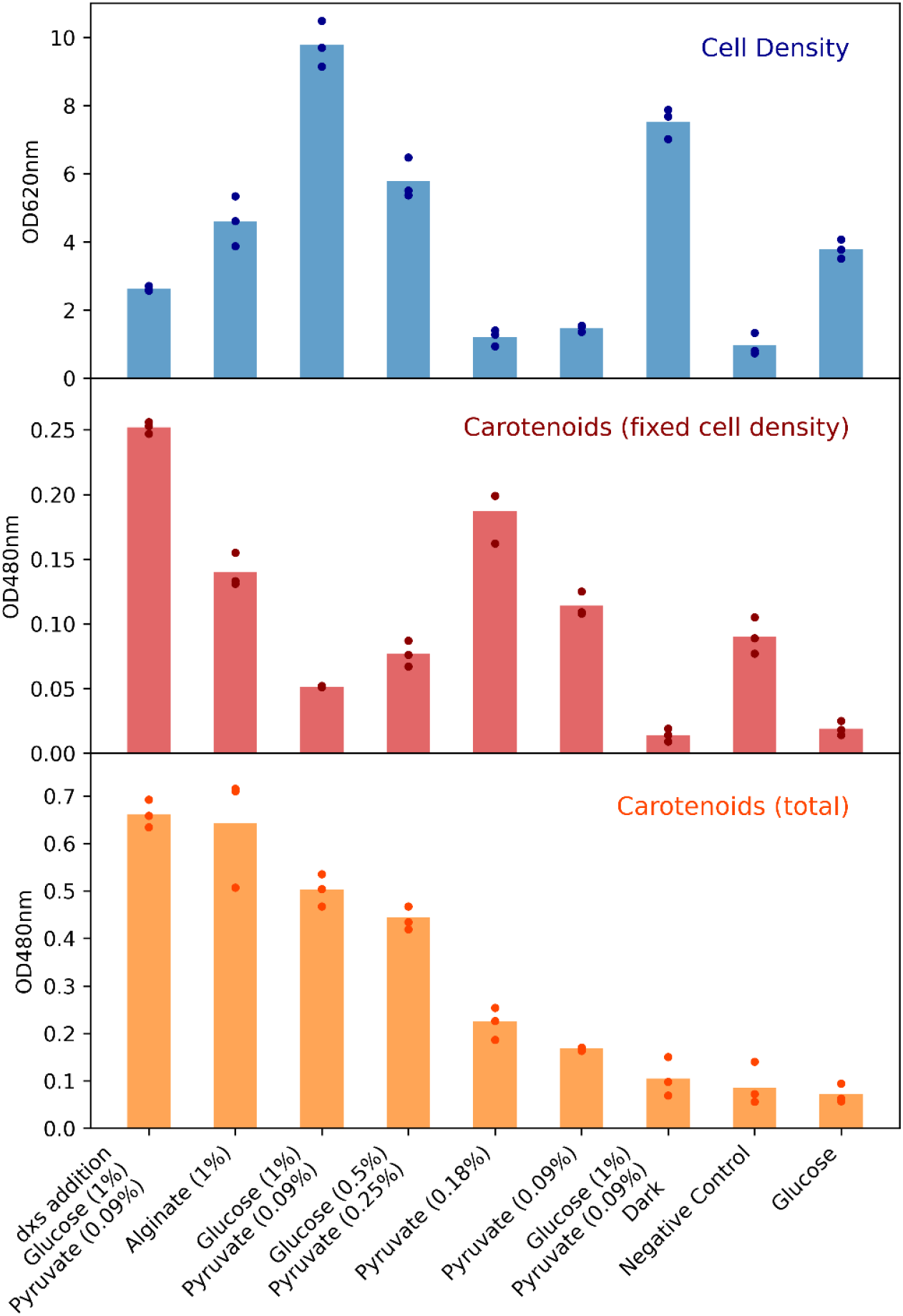
Cell density (OD620nm) and carotenoid (OD480nm) production following growth of *R. marinus* strain ISCaR-493 for 24 hours on glucose; mixture of glucose and pyruvate under light and dark conditions; alginate; pyruvate and without any carbon sources. Additionally, the modified strain TK-4 (Δ*trpB*Δ*purA::trpBdxs*_*T*.*thermophilus*_) was grown on a mixture of glucose and pyruvate. The carotenoids were always extracted from a fixed number of cells (1 mL of OD620nm = 1). The total amount of carotenoids was calculated by multiplying the cell density by the measured carotenoids. The average cell density of each culture condition is represented by a blue bar, the average measured carotenoid value as a red bar and the average total carotenoids by an orange bar. Dots represent individual replicates.

#### Glucose and pyruvate

*R. marinus* can grow on several monosaccharides, as predicted by the model. However, we have often observed better growth on oligo- and polysaccharides (data not shown). Growth of strain ISCaR-493 in defined medium with glucose (1%) as the sole carbon source resulted neither in high cell density nor high carotenoid production (Figure 4). *R. marinus* can utilize pyruvate as the sole carbon source (Table 2). Pyruvate is used in several pathways essential for growth and is the substrate, together with glyceraldehyde 3-phosphate, in the first step of the MEP terpenoid pathway (Figure 1 To increase both cell density and carotenoid production, pyruvate (0.09%) was added to the glucose-based medium. This resulted in increased carotenoid production and highly increased cell density (Figure 4). Visually, these cultures exhibited much stronger red color than the glucose cultures, which can be explained by both increased carotenoids yields and higher cell densities.

#### Impact of light

During the model reconstruction, a homolog for a light dependent regulatory gene was found in the carotenoid gene cluster in *R. marinus*. This indicates that carotenoid production in *R. marinus* is light induced (section 3.1). This was investigated here by cultivating ISCaR-493 in glucose (1%) and pyruvate (0.09%) medium in the dark and compared to the corresponding cultures grown in the light. The former cultures were colorless and the lack of carotenoids was confirmed by measurements (Figure 4).

#### Alginate

*R. marinus* can grow on many different polysaccharides (Table 2), making it an interesting candidate for processing 2^nd^ or 3^rd^ generation biomass, such as seaweed. Alginate is one of the major polysaccharides of brown algae. The products from alginate degradation are pyruvate and glyceraldehyde 3-phosphate (Figure 1), which are the same metabolites as used in the first step of the MEP terpenoid pathway. This raised the question whether *R. marinus* produces more carotenoids when grown on alginate, since it produces the two metabolites needed for the biosynthesis concurrently and in equal amounts. Cultivations in defined medium with alginate (1%) as the sole carbon source showed less cell density compared to glucose and pyruvate, but highly increased carotenoid production (Figure 4).

#### Glucose and pyruvate in equal quantities

To further examine if the availability of glyceraldehyde 3-phosphate and pyruvate in equal amounts results in higher carotenoid production, cultivation in defined medium with glucose (0.5%) and pyruvate (0,25%) was investigated. These cultivations showed lower cell density and higher carotenoid production compared to growth on glucose (1%) and pyruvate (0.09%) (Figure 4). The increased carotenoid production could be due to the equal availability of the two metabolites. Another possibility is that increased concentration of pyruvate alone in the medium caused higher carotenoid production.

#### Pyruvate

To examine if pyruvate alone affects the carotenoid production, two additional cultivations were set up, with pyruvate (0.09% and 0.18%) as the sole carbon source. The cell density in these cultures was low, only increased slightly after inoculation. This indicated that ISCaR-493 struggles to grow in liquid defined medium with pyruvate as the sole carbon source, which was surprising as growth was observed on agar medium (Table 2). The carotenoids per fixed cell density in the pyruvate cultures were much higher compared to cultures on glucose (1%) and pyruvate (0,09%). Additionally, increased pyruvate concentration resulted in increased carotenoid production (Figure 3). This suggests that the pyruvate is used for carotenoid production. Producing glyceraldehyde 3-phosphate from pyruvate costs energy (gluconeogenesis) and it cannot be determined from this data if this is the case for the observed growth. However, glycogen is an alternative source of glyceraldehyde 3-phosphate. The amount of glycogen in the biomass of *R. marinus* has been estimated as 14% [45] and is relatively high compared to other bacteria. Inclusion that could possibly contain glycogen can be discerned on electron micrographs of *R. marinus* [7]. Considering the natural habitat of *R. marinus* in coastal hot springs, it is not unreasonable to assume that it accumulates high levels of glycogen. Due to tides, the availability of nutrients in the surroundings of *R. marinus* varies widely and it is likely that the bacterium stores carbohydrates when they are in abundance in the environment. Since little or no growth was observed on pyruvate in liquid cultures it is likely that the cells experienced starvation and therefore started the breakdown of glycogen and carotenoid production. This was also seen for the negative control cultures without a carbon source (Figure 4). The cell density did not increase from inoculation, while the carotenoid production did.

#### Addition of the *dxs* gene from *T. thermophilus*

In an effort to increase carotenoid yields, the *dxs* gene from *T. thermophilus* was cloned on a shuttle vector into *R. marinus* strain SB-62 (ISCaR-493 derivative, Δ*trpB*Δ*purA*), resulting in the mutant strain TK-4 (Δ*trpB*Δ*purA::trpBdxs*_*T*.*thermophilus*_) (Supplementary file 5). The *dxs* gene encodes 1-deoxy-D-xylulose-5-phosphate synthase (DXS), which catalyzes the first step in the MEP terpenoid pathway (section 3.1) and could not be identified in the genomes of *R. marinus*. Compared to ISCaR-493, cultivation of TK-4 resulted in lower cell density but highly increased carotenoid production.

Presumably the added *dxs* gene resulted in a higher flux of carbons through the terpenoid and carotenoid pathways. However, it is also possible that this strain struggles to grow and responds by producing carotenoids. The dramatically lower cell density compared to ISCaR-493 can most likely be explained by the metabolic burden caused by the replication of the shuttle vector and the expression of its genes. Inserting the *dxs* gene in to the chromosome could reduce such effects.

In summary, these experiments showed that the highest cell density was obtained in glucose medium supplemented with pyruvate, while higher carotenoid production was observed during growth on alginate, with pyruvate added to a glucose-based medium and in the presence of light. It also showed that the carotenoid production per cell increased during starvation, indicating that yields can potentially be increased by either allowing the culture to reach and stay in stationary phase or transfer the cells after growth to new medium with limited or no carbon source. The motivation for the latter is that after the growth phase, the medium might not be optimal, e.g. due to accumulation of by-products that alter the pH, and the cells might stay alive and produce carotenoids longer in fresh medium. Finally, cloning the *dxs* gene from *T. thermophilus* in *R. marinus* resulted in the highest yields of carotenoids, but much lower cell density than the wild type strain ISCaR-493.

## 4. Conclusions

A manually curated genome-scale metabolic model of *R. marinus* DSM 4252^T^ was reconstructed and made publicly available (https://github.com/steinng/rmarinus). Experimental data from the literature and from this study was used to curate and validate the model. This includes growth data on various carbon sources, bioreactor cultivations and HPLC measurements of main metabolites, used for model validation, multiple studies on different metabolic pathways, components, genes and enzymes, and data on biomass components, which was used to formulate a species-specific biomass objective function.

The model was also evaluated for use with *R. marinus* ISCaR-493, from which the genetically modified SB-62 (Δ*trpB*Δ*purA*) was derived. The genome of strain ISCaR-493 was sequenced and the resulting draft genome was compared to that of strain DSM 4252^T^. This analysis showed that only seven model genes were absent in strain ISCaR-493 and four of them were replaced by genes encoding isozymes that exhibited high similarity to the DSM 4252^T^ enzymes. The remaining three genes are involved in EPS formation and xylan- and alginate degradation. EPSs of both strains have been previously studied [10] and shown to be of similar structures. It was also observed that strain ISCaR-493 grows well in defined medium with xylan and alginate as the sole carbon sources. In conclusion, this analysis suggests that the model is applicable for both strains DSM 4252^T^ and ISCaR-493. Both strains should be considered when any future changes or additions to the model reconstruction are made. Data on growth and metabolites was used to constrain the model and compare the experimental and simulated growth rates. This revealed that the model predicts correct growth rates for both strains.

Different aspects of the metabolism of *R. marinus* were reviewed during the reconstruction process. Here, an emphasis was on those with a potential biotechnological aspect, carotenoids in particular. Cell density and carotenoid production of strain ISCaR-493 grown at different conditions were investigated. Pyruvate addition to a glucose-based medium, highly increased cell density was obtained. Carotenoid production varied considerably under different growth conditions. Higher carotenoid yields were observed when pyruvate was present in the growth medium, alginate was used as the sole carbon source, cultivating the cells in light conditions and the cells experienced starvation. Additionally, we cloned the *dxs* gene from *T. thermophilus* on a shuttle vector into *R. marinus* and cultivation of the resulting mutant showed low cell density compared to ISCaR-493, but higher carotenoid production.

With its thermostable enzymes, wide range of potential carbon sources for growth and marketable products, *R. marinus* application potential is highly relevant in biotechnology and biorefineries. A genome-scale metabolic model helps us to understand its metabolism and should be useful in future strain designs.

## CRediT author statement

**Thordis Kristjansdottir**: Conceptualization, Methodology, Investigation, Writing – Original Draft, Writing – Review and Editing, Visualization. **Gudmundur O. Hreggvidsson:** Conceptualization, Investigation, Supervision, Funding acquisition, Writing – Original Draft, Writing – Review and Editing. **Sigmar Karl Stefansson:** Methodology, Investigation, Writing – Review and Editing, Visualization. **Elisabet Eik Gudmundsdottir:** Methodology, Investigation, Writing – Review and Editing. **Snaedis H. Bjornsdottir:** Methodology, Investigation, Writing – Review and Editing. **Olafur H. Fridjonsson:** Conceptualization, Writing – Review and Editing. **Eva Nordberg Karlsson:** Conceptualization, Writing – Review and Editing. **Justine Vanhalst:** Investigation, Writing – Review and Editing. **Birkir Reynisson:** Investigation, Writing – Review and Editing. **Steinn Gudmundsson:** Conceptualization, Supervision, Project administration, Funding acquisition, Writing – Review and Editing.

## Acknowledgements

The authors would like to thank Emanuel Y. C. Ron and Roya R. R. Sardari for the advice on how to cultivate *R. marinus* in bioreactors and to extract carotenoids, respectively, and Louna Maignien for assisting with the bioreactor cultivations.

