## Supplementary file 5 for "A genome-scale metabolic reconstruction provides insight into the metabolism of the thermophilic bacterium *Rhodothermus marinus*"

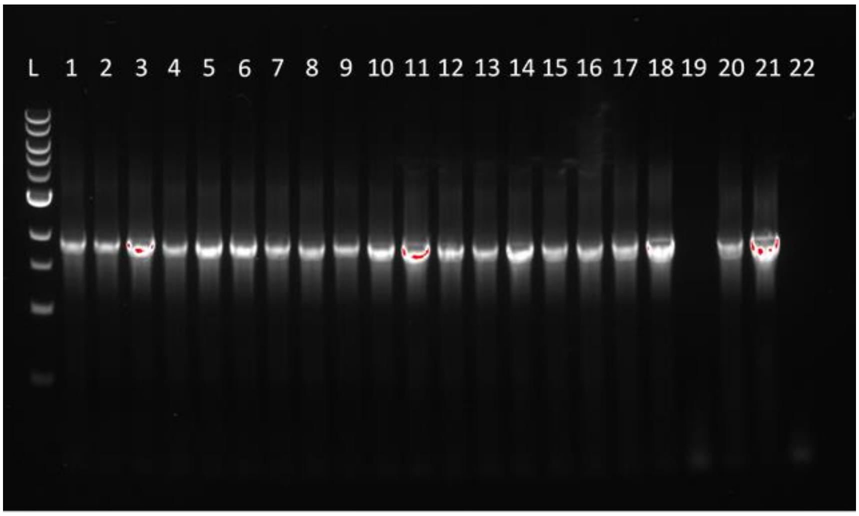


20 obtained *R. marinus dxs* clones picked for PCR (lanes 1-20). 19 of 20 were positive and 1 of 20 was negative. *T. thermophilus* gDNA (lane 21) was used as positive control and *R. marinus* SB-62 gDNA (lane 22) was used as negative control. 1kb ladder from BioLabs was used (lane L).
