## Supplementary file 2 for "A genome-scale metabolic reconstruction provides insight into the metabolism of the thermophilic bacterium *Rhodothermus marinus*"

### Biomass composition of *R. marinus*

The macromolecular composition of the biomass in *R. marinus* was obtained from *R. marinus* specific data found in literature [1]–[3], experimentally obtained in this study and when data on *R. marinus* was not available, experimental data for *E. coli* was used [4]. Amino acid [1], EPS [2], lipid [5], [6], carotenoid [3] and polyamine [7] compositions for *R. marinus* were available in literature. DNA and RNA compositions were calculated from the DSM 4252 genome.

| **Component** | | **Fraction (% w/w)** | **Molar mass (g mol$^{-1}$)** | **Coefficient (mmol gDW$^{-1}$)** | **Source** |
|---|---|---|---|---|---|
| Protein | PROT_RMAR_c | 45,8% | 128,0 | 3,5815 | *R. marinus* [1] |
| Glycogen | glycogen_c | 14,0% | 162,1 | 0,8643 | *R. marinus* [1] |
| Peptidoglycan | peptido_c | 2,50% | 991,0 | 0,0253 | *E. coli* [4] |
| EPS | EPS_RMAR_c | 0,36% | 1121,8 | 0,0032 | *R. marinus* [2] |
| LPS | lipidAds_c | 3,40% | 1323,7 | 0,0257 | *E. coli* [4] |
| DNA | DNA_RMAR | 2,50% | 304,2 | 0,0823 | *R. marinus* (This study) |
| RNA | RNA_RMAR | 13,0% | 322,7 | 0,4032 | *R. marinus* [1] |
| Lipids | LIP_RMAR_c | 16,2% | 970,6 | 0,1671 | *R. marinus* [1] |
| Carotenoids | CAROT_RMAR_c | 0,04% | 955,4 | 0,0004 | *R. marinus* [3] |
| Polyamines | POLAM_RMAR_c | 0,43% | 178,6 | 0,0241 | *E. coli* [4] |
| Soluble pool | SOL_c | 0,67% | | 1 | *E. coli* [4] |
| Inorganic ions | IONS_c | 1,00% | | 1 | *E. coli* [4] |
| | Total | 100% | | 3,5815 | |

**Biomass reaction:**

0.0004 CAROT_RMAR_c + 0.0822 DNA_RMAR_c + 0.0032 EPS_RMAR_c + IONS_c + 0.167 LIP_RMAR_c + 0.0241 POLAM_RMAR_c + 3.5814 PROT_RMAR_c + 0.4032 RNA_RMAR_c + SOL_c + 20.0 atp_c + 0.8643 glycogen_c + 20.0 h2o_c + 0.0257 lipidAds_c + 0.0252 peptido_c --> 20.0 adp_c + 20.0 h_c + 20.0 pi_c

#### Protein

Amino acid composition obtained mainly from *R. marinus* [1].

| **Amino acid** | **Fraction (mmol/g$_{protein}$)** | **Molar ratio** | **Molar mass (g mol$^{-1}$)** |
|---|---|---|---|
| Ala | 0,90 | 9,9% | 89,1 |
| Arg | 0,51* | 5,7% | 174,2 |
| Asx | 0,71 | 7,8% | |
| Asp | | 3,9% | 132,1 |
| Asn | | 3,9% | 132,1 |
| Cys | 0,16* | 1,8% | 121,2 |
| Glx | 1,09 | 12,1% | |
| Glu | | 6,0% | 146,6 |
| Gln | | 6,0% | 146,6 |
| Gly | 0,82 | 9,1% | 75,1 |

| | | | |
|---|---|---|---|
| His | 0,41* | 4,5% | 155,2 |
| Ile | 0,16 | 1,8% | 131,1 |
| Leu | 0,80 | 8,8% | 131,1 |
| Lys | 0,45 | 5,0% | 146,2 |
| Met | 0,20 | 2,2% | 149,2 |
| Phe | 0,31 | 3,4% | 165,2 |
| Pro | 0,53 | 5,8% | 115,1 |
| Ser | 0,47 | 5,2% | 105,1 |
| Thr | 0,54 | 6,0% | 119,1 |
| Trp | 0,10* | 1,1% | 204,2 |
| Tyr | 0,30 | 3,3% | 181,2 |
| Val | 0,62 | 6,8% | 117,2 |
| Total | 9,08 | 100% | 128,0 |

*Taken from *E. coli* [4]

**Protein reaction:**

0.0986 alatrna_c + 0.0565 argtrna_c + 0.0389 asntrna_c + 0.0389 asptrna_c + 0.306 atp_c + 0.0175 cystrna_c + 0.0603 glntrna_c + 0.0603 glutrna_c + 0.0908 glytrna_c + 2.0 gtp_c + 2.306 h2o_c + 0.0452 histrna_c + 0.0181 iletrna_c + 0.0876 leutrna_c + 0.0496 lystrna_c + 0.0217 mettrna_c + 0.034 phetrna_c + 0.0579 protrna_c + 0.0517 sertrna_c + 0.0599 thrtrna_c + 0.0109 trptrna_c + 0.0335 tyrtrna_c + 0.0682 valtrna_c --> PROT_RMAR_c + 0.306 adp_c + 2.0 gdp_c + 2.306 h_c + 2.306 pi_c + 0.0986 trnaala_c + 0.0565 trnaarg_c + 0.0389 trnaasn_c + 0.0389 trnaasp_c + 0.0175 trnacys_c + 0.1206 trnaglu_c + 0.0908 trnagly_c + 0.0452 trnahis_c + 0.0181 trnaile_c + 0.0876 trnaleu_c + 0.0496 trnalys_c + 0.0217 trnamet_c + 0.034 trnaphe_c + 0.0579 trnapro_c + 0.0517 trnaser_c + 0.0599 trnathr_c + 0.0109 trnatrp_c + 0.0335 trnatyr_c + 0.0682 trnaval_c

### DNA

The DNA fraction of the biomass was estimated in this study. A dsDNA stock solution was made by diluting 102,7g/ml (ABS @ 260nm using Nanodrop) λ-dsDNA solution x50, giving 2,1 g/ml.

Standard curve measurements of PicoGreen DNA assay using λ-dsDNA standard

| Part Stock Solution | Conc. g⁄l | ABS | | | Mean | Mean-blank |
|---|---|---|---|---|---|---|
| 0 | 0 | 120 | 106 | 136 | 121 | 0 |
| 0,001 | 0,002 | 958 | 1016 | 1166 | 1047 | 926 |
| 0,01 | 0,021 | 3808 | 3958 | 3454 | 3740 | 3619 |
| 0,1 | 0,205 | 32942 | 32990 | 31786 | 32573 | 32452 |
| 0,5 | 1,027 | 162135 | 167970 | 171127 | 167077 | 166957 |
| 1 | 2,054 | 346972 | 370334 | 332978 | 350095 | 349974 |

$y = 170000x – 1500$
$r^2 = 0;999$

Lysed *R. marinus* cells measured

| ABS | | | Mean | Mean-blank |
|---|---|---|---|---|
| 41341 | 42907 | 42439 | 42229 | 42108 |

DNA ratio calculated from DSM 4252 genome

| Nucleotide | Molar ratio | Molar mass (g mol$^{-1}$) |
|---|---|---|
| dAMP | 17,8% | 331,2 |
| dCMP | 32,2% | 289,2 |
| dGMP | 32,2% | 304,2 |
| dTMP | 17,8% | 304,2 |
| Total | 100% | 304,2 |

**DNA reaction:**

1.37 atp_c + 0.178 datp_c + 0.322 dctp_c + 0.322 dgtp_c + 0.178 dttp_c --> DNA_RMAR_c + 1.37 adp_c + 1.37 h_c + 1.37 pi_c + ppi_c

### RNA

Calculated from DSM 4252 genome, assuming equal transcription

| Nucleotide | Molar ratio | Molar mass (g mol$^{-1}$) |
|---|---|---|
| CMP | 32,2% | 305,2 |
| GMP | 33,0% | 345,2 |
| UMP | 17,6% | 306,2 |
| AMP | 17,2% | 329,2 |
| Total | 100% | 322,7 |

**RNA reaction:**

0.572 atp_c + 0.322 ctp_c + 0.33 gtp_c + 0.176 utp_c --> RNA_RMAR_c + 0.4 adp_c + 0.4 h_c + 0.4 pi_c + ppi_c

### Lipids

Fatty acid composition in *R. marinus* obtained from [6]

| Fatty acid | Molar ratio |
|---|---|
| iC14 | 1,6% |
| nC14 | 0,2% |
| iC15 | 8,4% |
| aC15 | 19,1% |
| nC15 | 0,2% |
| iC16 | 13,0% |
| nC16 | 5,4% |
| iC17 | 26,4% |
| aC17 | 18,4% |
| nC17 | 0,7% |
| iC18 | 4,7% |
| nC18 | 0,9% |

| | |
|---|---|
| iC19 | 0,2% |
| aC19 | 0,4% |

Phospholipid composition in *R. marinus* obtained from [5]

| **Lipid** | **Molar ratio** |
|---|---|
| PE | 50,0% |
| DPG | 40,0% |
| PG | 10,0% |

Taken together:

| **Lipid** | **Molar ratio** | **Molar mass** |
|---|---|---|
| pgi14 | 0,16% | 666,9 |
| pg140 | 0,02% | 666,9 |
| pgi15 | 0,84% | 694,9 |
| pga15 | 1,92% | 694,9 |
| pg150 | 0,02% | 694,9 |
| pgi16 | 1,31% | 723,0 |
| pg160 | 0,54% | 723,0 |
| pgi17 | 2,65% | 751,0 |
| pga17 | 1,85% | 751,0 |
| pg170 | 0,07% | 751,0 |
| pgi18 | 0,47% | 779,1 |
| pg180 | 0,09% | 779,1 |
| pgi19 | 0,02% | 807,1 |
| pga19 | 0,04% | 807,1 |
| pei14 | 0,80% | 635,9 |
| pe140 | 0,10% | 635,9 |
| pei15 | 4,22% | 663,9 |
| pea15 | 9,59% | 663,9 |
| pe150 | 0,10% | 663,9 |
| pei16 | 6,53% | 692,0 |
| pe160 | 2,71% | 692,0 |

| **Lipid** | **Molar ratio** | **Molar mass** |
|---|---|---|
| pei17 | 13,25% | 720,0 |
| pea17 | 9,24% | 720,0 |
| pe170 | 0,35% | 720,0 |
| pei18 | 2,36% | 748,1 |
| pe180 | 0,45% | 748,1 |
| pei19 | 0,10% | 776,1 |
| pea19 | 0,20% | 776,1 |
| dpgi14 | 0,64% | 1241,7 |
| dpg140 | 0,08% | 1241,7 |
| dpgi15 | 3,37% | 1297,8 |
| dpga15 | 7,67% | 1297,8 |
| dpg150 | 0,08% | 1297,8 |
| dpgi16 | 5,22% | 1353,9 |
| dpg160 | 2,17% | 1353,9 |
| dpgi17 | 10,60% | 1410,0 |
| dpga17 | 7,39% | 1410,0 |
| dpg170 | 0,28% | 1410,0 |
| dpgi18 | 1,89% | 1466,1 |
| dpg180 | 0,36% | 1466,1 |
| dpgi19 | 0,08% | 1522,2 |
| dpga19 | 0,16% | 1522,2 |
| Total | 100% | 970,6 |

**Lipid reaction:**

0.0008 dpg140_c + 0.0008 dpg150_c + 0.0217 dpg160_c + 0.0028 dpg170_c + 0.0036 dpg180_c + 0.0767 dpga15_c + 0.0739 dpga17_c + 0.0016 dpga19_c + 0.0064 dpgi14_c + 0.0337 dpgi15_c + 0.0522 dpgi16_c + 0.106 dpgi17_c + 0.0189 dpgi18_c + 0.0008 dpgi19_c + 0.001 pe140_c + 0.001 pe150_c + 0.0271 pe160_c + 0.0035 pe170_c + 0.0045 pe180_c + 0.0959 pea15_c + 0.0924 pea17_c + 0.002 pea19_c + 0.008 pei14_c + 0.0422 pei15_c + 0.0653 pei16_c + 0.1325 pei17_c + 0.0236 pei18_c + 0.001 pei19_c + 0.0002 pg140_c + 0.0002 pg150_c + 0.0054 pg160_c + 0.0007 pg170_c + 0.0009 pg180_c + 0.0192 pga15_c + 0.0185 pga17_c + 0.0004 pga19_c + 0.0016 pgi14_c + 0.0084 pgi15_c + 0.0131 pgi16_c + 0.0265 pgi17_c + 0.0047 pgi18_c + 0.0002 pgi19_c --> LIP_RMAR_c

### Carotenoids

Carotenoid composition in *R. marinus* obtained from [3]

| **Carotenoid** | | **Molar ratio** | **Molar mass (g mol$^{-1}$)** |
|---|---|---|---|
| hglpyrcarote | Non-acetylated | 3,30% | 745,0 |
| ddcahglpyrcarote | Iso-C12:0 | 5,08% | 927,3 |
| myrshglpyrcarote | n-C12:0 | 0,56% | 955,4 |
| palmhglpyrcarote | Iso-C13:0 | 7,34% | 983,4 |
| odechglpyrcarote | Anteiso-C13:0 | 9,03% | 1011,5 |
| pdechglpyrcarote | Iso-C14:0 | 4,51% | 969,4 |
| 10mudechglpyrcarote | n-C14:0 | 2,26% | 927,3 |
| 12mtdechglpyrcarote | Iso-C15:0 | 5,64% | 955,4 |
| 14mpdechglpyrcarote | Anteiso-C15:0 | 10,72% | 983,4 |
| 11mddechglpyrcarote | n-C15:0 | 0,56% | 941,3 |
| 13mmyrshglpyrcarote | Iso-C16:0 | 3,95% | 969,4 |
| 15mpalmhglpyrcarote | n-C16:0 | 3,39% | 997,5 |
| 10mddechglpyrcarote | Iso-C17:0 | 0,56% | 941,3 |
| 12mmyrshglpyrcarote | Anteiso-C17:0 | 1,69% | 969,4 |
| glpyrcarote | Non-acetylated | 2,50% | 729,0 |
| ddcaglpyrcarote | Iso-C12:0 | 3,50% | 911,3 |
| myrsglpyrcarote | n-C12:0 | 0,39% | 939,4 |
| palmglpyrcarote | Iso-C13:0 | 6,22% | 967,4 |
| odecglpyrcarote | Anteiso-C13:0 | 7,00% | 995,5 |
| pdecglpyrcarote | Iso-C14:0 | 2,33% | 953,4 |
| 10mudecglpyrcarote | n-C14:0 | 1,56% | 911,3 |
| 12mtdecglpyrcarote | Iso-C15:0 | 3,89% | 939,4 |
| 14mpdecglpyrcarote | Anteiso-C15:0 | 7,39% | 967,4 |
| 11mddecglpyrcarote | n-C15:0 | 0,39% | 925,3 |
| 13mmyrsglpyrcarote | Iso-C16:0 | 1,95% | 953,4 |
| 15mpalmglpyrcarote | n-C16:0 | 2,33% | 981,5 |
| 10mddecglpyrcarote | Iso-C17:0 | 0,39% | 925,3 |
| 12mmyrsglpyrcarote | Anteiso-C17:0 | 1,17% | 953,4 |
| 14mpalmglpyrcarote | n-C18:0 | 0,39% | 981,5 |
| Total | | 100% | 955,4 |

**Carotenoid reaction:**

0.004 10mddecglpyrcarote_c + 0.006 10mddechglpyrcarote_c + 0.016 10mudecglpyrcarote_c + 0.023 10mudechglpyrcarote_c + 0.004 11mddecglpyrcarote_c + 0.006 11mddechglpyrcarote_c + 0.012 12mmyrsglpyrcarote_c + 0.017 12mmyrshglpyrcarote_c + 0.039 12mtdecglpyrcarote_c + 0.056 12mtdechglpyrcarote_c + 0.019 13mmyrsglpyrcarote_c + 0.04 13mmyrshglpyrcarote_c + 0.004 14mpalmglpyrcarote_c + 0.074 14mpdecglpyrcarote_c + 0.107 14mpdechglpyrcarote_c + 0.023 15mpalmglpyrcarote_c + 0.034 15mpalmhglpyrcarote_c + 0.035 ddcaglpyrcarote_c + 0.051 ddcahglpyrcarote_c + 0.025 glpyrcarote_c + 0.033 hglpyrcarote_c + 0.004 myrsglpyrcarote_c + 0.006 myrshglpyrcarote_c + 0.07 odecglpyrcarote_c + 0.09 odechglpyrcarote_c + 0.062 palmglpyrcarote_c

+ 0.073 palmhglpyrcarote_c + 0.023 pdecglpyrcarote_c + 0.045 pdechglpyrcarote_c --> CAROT_RMAR_c

### Polyamine

Polyamine composition in *R. marinus* obtained from [7]

| **Polyamines** | **Fraction (mmol $gDW^{-1}$)** | **Molar ratio** | **Molar mass (g $mol^{-1}$)** |
|---|---:|---:|---:|
| Putrescine | 0,000245 | 1,02% | 90,2 |
| Cadaverine | 0,000245 | 1,02% | 104,2 |
| Spermidine | 0,012840 | 53,30% | 148,3 |
| Spermine | 0,007337 | 30,46% | 206,4 |
| Thermopentamine | 0,000978 | 4,06% | 261,5 |
| N4-Aminopropylspermidine | 0,000611 | 2,54% | 206,4 |
| N4-bis(aminopropyl)spermidine | 0,001834 | 7,61% | 247,5 |
| Total | 0,024091 | 100% | 178,6 |

### Exopolysaccharides

Exopolysaccharide composition in *R. marinus* obtained from [2]

| **EPS unit** | **Molar ratio** | **Molar mass (minus udp/dtdp) (g $mol^{-1}$)** |
|---|---:|---:|
| UDP-glucose | 0,5 | 160,125 |
| dTDP-glucose | 0,5 | 163,149 |
| UDP-arabinose | 2,45 | 130,099 |
| UDP-xylose | 4,93 | 130,099 |
| Total | | 1121,768 |

**EPS reaction:**

0.5 dtdpglu_c + h2o_c + 2.45 udparab_c + 0.5 udpg_c + 4.93 udpxyl_c --> EPS_RMAR_c + 0.5 dtdp_c + 7.88 udp_c

### Vitamins, cofactors and inorganic ions

Soluble pool and inorganic ions adapted from *E. coli* [4]

**Soluble pool reaction:**

0.00022 10fthf_c + 0.00022 5mthf_c + 0.00028 accoa_c + 0.00022 adocbl_c + 0.00022 btn_c + 0.00022 chor_c + 0.00017 coa_c + 0.00022 fad_c + 0.00022 gthrd_c + 0.00022 hemeA_c + 0.00022 hemeO_c + 3e-05 malcoa_c + 0.00022 mlthf_c + 0.00179 nad_c + 4e-05 nadh_c + 0.00011 nadp_c + 0.00034 nadph_c + 0.00022 pheme_c + 0.00022 pydx5p_c + 0.00022 ribflv_c + 0.00022 sheme_c + 0.0001 succoa_c + 0.00022 thf_c + 6e-05 udcpdp_c --> SOL_c

**Inorganic ions reaction:**

0.0045 ca2_c + 0.003 cobalt2_c + 0.003 cu2_c + 0.0068 fe2_c + 0.0068 fe3_c + 0.1704 k_c + 0.0076 mg2_c + 0.003 mn2_c + 0.003 mobd_c + 0.0038 na1_c + 0.0114 nh4_c + 0.0038 pi_c + 0.0038 so4_c + 0.003 zn2_c --> IONS_c

**GAM and NGAM**

No data for energy requirements in *R. marinus* is available. NGAM (non-growth associated maintenance) was set to 1 mmol/gDW/h but GAM (growth associated maintenance) was estimated by constraining the model with experimental uptake- and secretion rates of glucose, pyruvate, lactate and acetate, and growth rate of *R. marinus*. The GAM was estimated to be 20 mmol/gDW/h. This is relatively low compared to many other models, which can be explained by the energy accounted for in biosynthetic reactions for the macromolecules. The dataset used for this estimation (supplementary file 1) was not used again for validation.

### Sensitivity analysis

The sensitivity of predicted growth rate to variation in biomass and energy components was investigated. The coefficient of each component was varied by 50% at different glucose uptake rates while the growth rate was predicted. Predicted growth rate was most sensitive to changes in the protein component, followed by the lipid component.

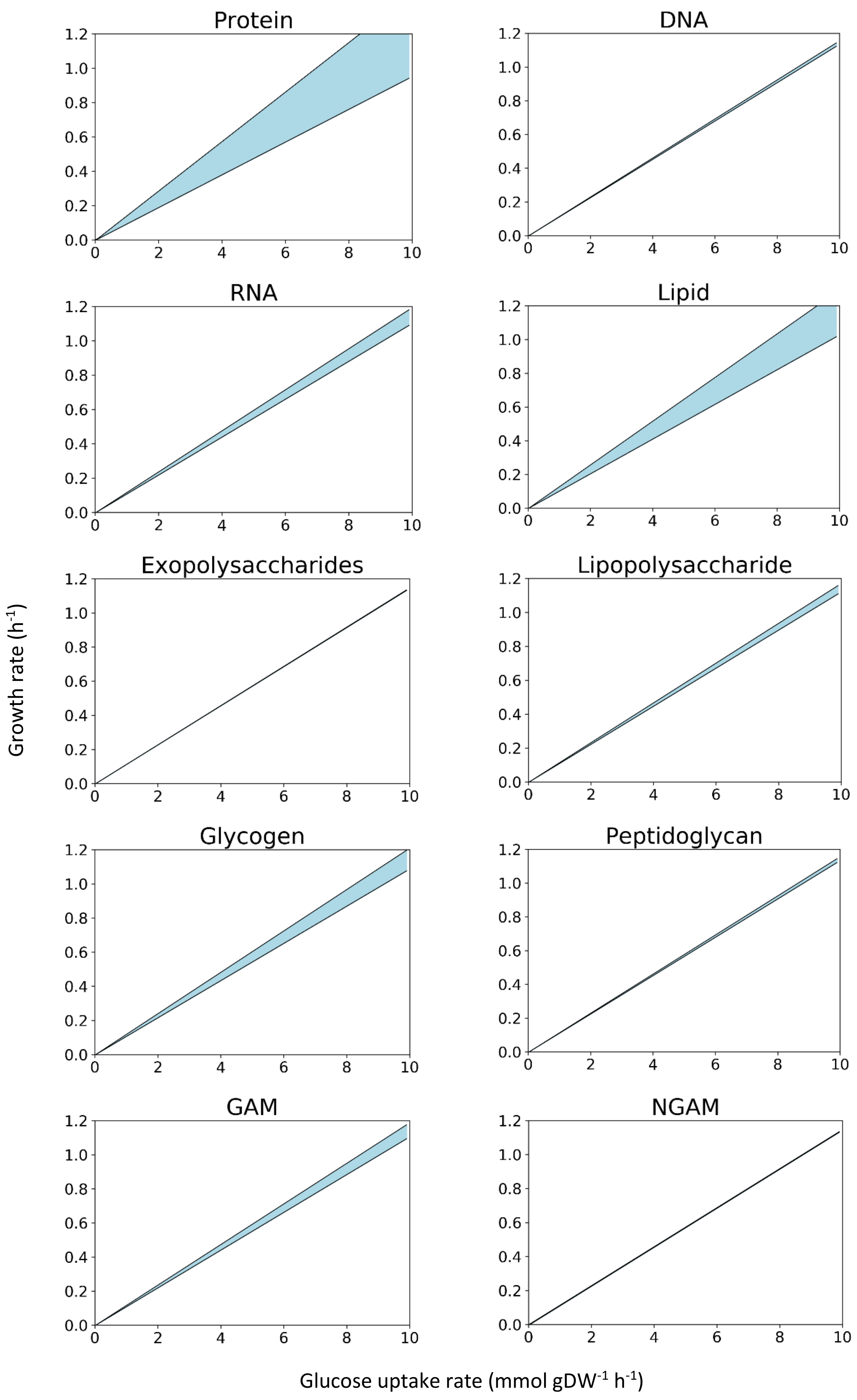

Protein
DNA
RNA
Lipid
Exopolysaccharides
Lipopolysaccharide
Glycogen
Peptidoglycan
GAM
NGAM
Growth rate (h-1)
Glucose uptake rate (mmol gDW-1 h-1)
