## Supplementary file 3 for "A genome-scale metabolic reconstruction provides insight into the metabolism of the thermophilic bacterium *Rhodothermus marinus*"

Rmarinus_578 Expand All Readme 2021-01-07 17:06

### Independent Section

Contains tests that are independent of the class of modeled organism, a model's complexity or types of identifiers that are used to describe its components.

#### Consistency

| | | |
|---|---|---|
| Stoichiometric Consistency | 100.0% | X3 |
| Mass Balance | 98.8% | |
| Charge Balance | 98.6% | |
| Metabolite Connectivity | 100.0% | |
| Unbounded Flux In Default Medium | 98.6% | |
| Sub Total | 99% | X3 |

#### Annotation - Metabolites

| | |
|---|---|
| Presence of Metabolite Annotation | 68.3% |
| Metabolite Annotations Per Database | Info |
| pubchem.compound | 0.0% |
| kegg.compound | 56.4% |
| seed.compound | 58.1% |
| inchikey | 0.0% |
| inchi | 0.0% |
| chebi | 56.9% |
| hmdb | 38.1% |
| reactome | 0.0% |
| metanetx.chemical | 60.4% |
| bigg.metabolite | 68.3% |
| biocyc | 52.0% |
| Metabolite Annotation Conformity Per Database | Info |
| pubchem.compound | 0.0% |
| kegg.compound | 100.0% |
| seed.compound | 100.0% |
| inchikey | 0.0% |
| inchi | 0.0% |
| chebi | 100.0% |
| hmdb | 100.0% |
| reactome | 0.0% |
| metanetx.chemical | 100.0% |
| bigg.metabolite | 98.2% |
| biocyc | 100.0% |
| Uniform Metabolite Identifier Namespace | 100.0% |

### Specific Section

Covers general statistics and specific aspects of a metabolic network that are not universally applicable. See readme for more details.

#### SBML

| | |
|---|---|
| SBML Level and Version | Errored |
| FBC enabled | Errored |

#### Basic Information

| | |
|---|---|
| Model Identifier | Rmarinus_578 |
| Total Metabolites | 871 |
| Total Reactions | 929 |
| Total Genes | 578 |
| Total Compartments | 3 |
| Metabolic Coverage | 1.61 |

#### Metabolite Information

| | |
|---|---|
| Unique Metabolites | 784 |
| Duplicate Metabolites in Identical Compartments | 0 |
| Metabolites without Charge | 0 |
| Metabolites without Formula | 0 |
| Medium Components | 22 |

#### Reaction Information

| | |
|---|---|
| Purely Metabolic Reactions | 789 |
| Purely Metabolic Reactions with Constraints | 1 |
| Transport Reactions | 92 |
| Transport Reactions with Constraints | 0 |
| Thermodynamic Reversibility of Purely Metabolic Reactions | 0.33 |
| Reactions With Partially Identical Annotations | 0.02 |
| Duplicate Reactions | 0.00 |
| Reactions With Identical Genes | 0.47 |

#### Gene-Protein-Reaction (GPR) Associations

| | |
|---|---|
| Reactions without GPR | 158 |
| Fraction of Transport Reactions without GPR | 0.59 |
| Enzyme Complexes | 70 |

Rmarinus_578 Expand All Readme 2021-01-07 17:06

#### Annotation - Reactions

| | |
|---|---|
| Presence of Reaction Annotation | 91.1% |
| Reaction Annotations Per Database | Info |
| rhea | 30.2% |
| kegg.reaction | 28.5% |
| seed.reaction | 0.0% |
| metanetx.reaction | 42.4% |
| bigg.reaction | 45.6% |
| reactome | 0.0% |
| ec-code | 81.6% |
| brenda | 0.0% |
| biocyc | 30.1% |
| Reaction Annotation Conformity Per Database | Info |
| rhea | 99.7% |
| kegg.reaction | 100.0% |
| seed.reaction | 0.0% |
| metanetx.reaction | 100.0% |
| bigg.reaction | 100.0% |
| reactome | 0.0% |
| ec-code | 99.2% |
| brenda | 0.0% |
| biocyc | 100.0% |
| Uniform Reaction Identifier Namespace | 100.0% |
| Sub Total | 72% |

#### Annotation - Genes

| | |
|---|---|
| Presence of Gene Annotation | 99.8% |
| Gene Annotations Per Database | Info |
| refseq | 0.0% |
| uniprot | 0.0% |
| ecogene | 0.0% |
| kegg.genes | 0.0% |
| ncbigi | 0.0% |
| ncbigene | 0.0% |
| ncbiprotein | 99.7% |
| ccds | 0.0% |
| hprd | 0.0% |
| asap | 0.0% |
| Gene Annotation Conformity Per Database | Info |
| refseq | 0.0% |

| | |
|---|---|
| Biomass Consistency | 0.2[illegible] |
| Biomass Production In Default Medium | 0.2[illegible] |
| Unrealistic Growth Rate In Default Medium | fals[illegible] |
| Biomass Production In Complete Medium | 41.0 |
| Blocked Biomass Precursors In Default Medium | 0 |
| Blocked Biomass Precursors In Complete Medium | 0 |
| Ratio of Direct Metabolites in Biomass Reaction | 0.0[illegible] |
| Number of Missing Essential Biomass Precursors | 35 |

#### Energy Metabolism

| | |
|---|---|
| Non-Growth Associated Maintenance Reaction | 1 |
| Growth-associated Maintenance in Biomass Reaction | true |
| Number of Reversible Oxygen-Containing Reactions | 4 |
| Erroneous Energy-generating Cycles | Info |
| MNXM3 | Skipped |
| MNXM63 | Skipped |
| MNXM51 | Skipped |
| MNXM121 | Skipped |
| MNXM423 | Skipped |
| MNXM6 | Skipped |
| MNXM10 | Skipped |
| MNXM38 | Skipped |
| MNXM208 | Skipped |
| MNXM191 | Skipped |
| MNXM223 | Skipped |
| MNXM7517 | Skipped |
| MNXM12233 | Skipped |
| MNXM558 | Skipped |
| MNXM21 | Skipped |
| MNXM89557 | Skipped |

#### Network Topology

| | |
|---|---|
| Universally Blocked Reactions | 53 |
| Orphan Metabolites | 5 |
| Dead-end Metabolites | 14 |
| Stoichiometrically Balanced Cycles | 18 |
| Metabolite Production In Complete Medium | 222 |
| Metabolite Consumption In Complete Medium | 239 |

#### Matrix Conditioning

Rmarinus_578 Expand All Readme 2021-01-07 17:06

| | | |
|---|---|---|
| ecogene | 0.0% | ⌄ |
| kegg.genes | 0.0% | ⌄ |
| ncbigi | 0.0% | ⌄ |
| ncbigene | 0.0% | ⌄ |
| ncbiprotein | 100.0% | ⌄ |
| ccds | 0.0% | ⌄ |
| hprd | 0.0% | ⌄ |
| asap | 0.0% | ⌄ |
| Sub Total | 40% | ⌄ |

#### Annotation - SBO Terms

| | | |
|---|---|---|
| Metabolite General SBO Presence | 0.0% | ⌄ |
| Metabolite SBO:0000247 Presence | 0.0% | ⌄ |
| Reaction General SBO Presence | 45.6% | ⌄ |
| Metabolic Reaction SBO:0000176 Presence | 0.0% | ⌄ |
| Transport Reaction SBO:0000185 Presence | 0.0% | ⌄ |
| Exchange Reaction SBO:0000627 Presence | 68.2% | ⌄ |
| Demand Reaction SBO:0000628 Presence | 0.0% | ⌄ |
| Sink Reactions SBO:0000632 Presence | Skipped | ⌄ |
| Gene General SBO Presence | 0.0% | ⌄ |
| Gene SBO:0000243 Presence | 0.0% | ⌄ |
| Biomass Reactions SBO:0000629 Presence | 0.0% | ⌄ |
| Sub Total | 10% | X2 ⌄ |

| | | |
|---|---|---|
| Total Score | 55% | ⌄ |

Total Score

# 55%

Score per Category

| | | |
|---|---|---|
| [illegible] Relations | 31 | ⌄ |
| Rank | 840 | ⌄ |
| Degrees Of Freedom | 89 | ⌄ |

#### Experimental Data Comparison

| | | |
|---|---|---|
| Growth Prediction | Skipped | ⌄ |
| Gene Essentiality Prediction | Skipped | ⌄ |

#### Misc. Tests

#### Environment

| | |
|---|---|
| Python Version | 3.6.12 |
| Platform | Linux |
| Memote Version | 0.11.1 |

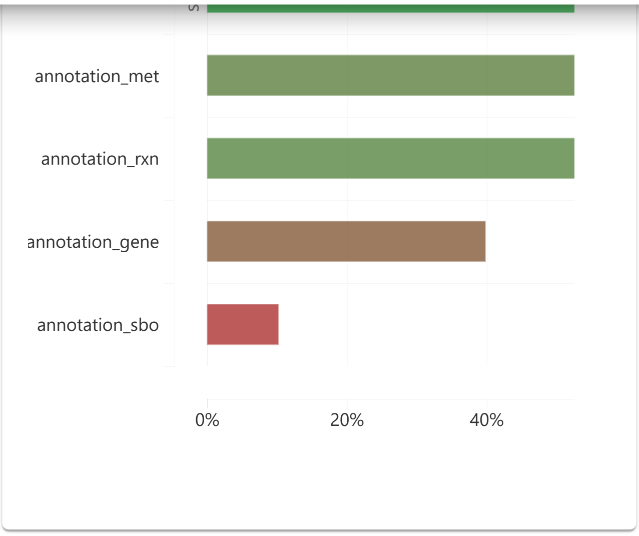
annotation_met
annotation_rxn
annotation_gene
annotation_sbo
0%
20%
40%
